# Effects Of Lymphocyte Trafficking on Antiretroviral Drug Distribution within the Lymph Node

**DOI:** 10.64898/2026.10.28.762578

**Authors:** Sobia Anam, Michael J. Donzanti, Allyson M. Dang, Sasan Paryad-Zanjani, Michael J. Piovoso, Jason P. Gleghorn, Ryan Zurakowski

## Abstract

The lymph node (LN) lobule is physically isolated by selective transport barriers, including high endothelial venules and sinusoidal endothelium, that restrict free diffusion of many antiretroviral (ARV) drugs used to treat HIV, especially non-lipophilic compounds. Recent spatial pharmacokinetic analyses have suggested that passive diffusion alone cannot explain observed ARV concentrations in lymphoid tissues, raising the possibility that lymphocytes may act as incidental drug carriers. To investigate the relative contributions of free and cell-mediated drug trafficking, we conducted murine experiments in which drug was delivered intravenously to a murine model, with lymphocyte entry into the lobule blocked by a functional antibody. The experimental results show that blocking lymphocyte entry into the LN increases raw intracellular drug concentration within the LN for tenofovir and efavirenz, but not raltegravir. To understand the mechanism behind this observation, we developed a computational transport model. The model consists of a mechanistic physiologically based pharmacokinetic framework with four compartments: plasma, peripheral blood mononuclear cells, LN extracellular fluid, and LN intracellular space, while dynamically accounting for changes in LN extracellular fluid volume driven by lymphocyte trafficking. Our simulations predicted that blocking lymphocyte trafficking increases extracellular fluid volume and, counterintuitively, elevates intracellular drug concentration within the LN, with the effect size depending on the drug’s intracellular equilibration kinetics. Raltegravir, which equilibrates rapidly between extracellular and intracellular compartments, is insensitive to this mechanism, a model prediction confirmed by the experimental null result. In conclusion, our study identifies extracellular fluid volume dynamics within the lymph node as a previously unrecognized, drug class-dependent modulator of intranodal ARV distribution. This mechanism operates independently of barrier-level penetration and has direct implications for drug efficacy in patients with altered immune trafficking.

**Study Highlights:** *CURRENT KNOWLEDGE ON THE TOPIC:* Combination antiretroviral therapy (cART) effectively suppresses plasma HIV levels but does not cure the infection, in part due to the presence of pharmacological sanctuary sites, such as lymph nodes (LNs). The transport of antiretrovirals (ARVs) into LNs is restricted by endothelial barriers and sinusoidal endothelium, and drug entry may occur through multiple pathways, including passive diffusion, active transport, and cell-mediated transport. However, the exact mechanisms of ARV distribution within lymph node (LN) lobules remain poorly understood.

*QUESTION ADDRESSED BY THIS STUDY:* This study investigates the role of lymphocyte trafficking in transporting ARVs into the LN and how modulating lymphocyte entry affects intracellular drug concentrations within the LN.

*WHAT THIS STUDY ADDS TO OUR KNOWLEDGE:* Blocking lymphocyte entry into the LN paradoxically increases intracellular ARV concentration for tenofovir (TFV) and efavirenz (EFV), but not raltegravir (RAL). A mechanistic multi-compartment physiologically based pharmacokinetic model incorporating extracellular fluid (ECF) volume dynamics within the fixed LN volume explains this finding: when fewer lymphocytes occupy the LN, ECF volume expands, drug gradients steepen, and intracellular uptake increases. This identifies ECF volume dynamics as a previously unrecognized modulator of intranodal drug distribution, operating independently of barrier-level penetration.

*POTENTIAL IMPACT ON DRUG DISCOVERY, DEVELOPMENT, AND THERAPEUTICS:* This work may help identify mechanisms that limit antiretroviral exposure within lymph nodes and improve understanding of how lymphocyte trafficking contributes to drug distribution within lymphoid tissue. These insights could support the optimization of drug properties, dosing strategies, and delivery approaches for therapies that require effective lymphoid tissue exposure.

## INTRODUCTION

Despite being one of the most studied viruses in history, HIV continues to evade curative treatment, persisting in hidden reservoirs even after decades of medical advancement. While antiretroviral therapy (ART) has transformed HIV from a fatal diagnosis into a manageable condition, it still cannot fully eradicate the virus^1–4^. A critical obstacle is that antiretroviral (ARV) drug concentrations within lymph node (LN) tissues are substantially lower than in peripheral blood. This deficit correlates directly with ongoing HIV replication in lymphoid tissues even during suppressive therapy^5^. Several studies have shown that many ARV drugs exhibit limited penetration into the lymph nodes (LNs) of treated individuals, a phenomenon associated with elevated HIV RNA levels in lymphoid tissues^6^. These pharmacological sanctuary sites represent one of the most significant remaining barriers to HIV cure^6,7^.

The LN lobule is structurally isolated from the bloodstream by highly selective transport barriers, including high endothelial venules (HEVs) and sinusoidal endothelium, that substantially restrict drug access and create conditions that allow a sanctuary to form^6,8–16^. Quantitative modeling has established that this barrier-level penetration failure has direct clinical consequences: limited intranodal drug exposure is sufficient to drive virological failure and resistance emergence^17^. Yet even accounting for these barriers, a critical disconnect remains between predicted and observed ARV distributions in lymphoid tissues. Diffusion-based models fitted to spatial mass spectrometry data consistently yield diffusivity estimates that are too low to be explained by free diffusion, especially for non-lipophilic drugs such as tenofovir and atazanavir^18^. These values, instead, align more closely with known lymphocyte motility parameters, suggesting that passive diffusion cannot fully account for drug penetration into the LN lobule. Given the structural isolation of the LN lobule behind the HEV^19^, these discrepancies point compellingly to an additional, unrecognized transport mechanism governing intranodal drug distribution.

The high rate of lymphocyte trafficking through LNs, with millions of cells entering and exiting via HEVs continuously^20^, raises the possibility that lymphocytes serve as incidental ARV carriers, particularly for non-lipophilic species. Yet, existing ARV pharmacokinetic (PK) models do not explicitly account for lymphocyte-mediated drug transport and how drug exposure is affected when lymphocyte trafficking is altered. This limits their utility for predicting therapeutic efficacy under conditions such as inflammation or immunosuppression^21–24^. To directly test this lymphocyte carrier hypothesis, we employed an L-selectin functional antibody blocking (FAB) approach to selectively prevent lymphocyte entry into the LN lobule in a murine model, while leaving passive diffusion and other transport pathways intact^25–28^.

Contrary to the carrier hypothesis, blocking lymphocyte entry paradoxically *increased* intracellular ARV concentration within the LN. To understand this counterintuitive result, we developed a mechanistic multi-compartment physiologically based pharmacokinetic (PBPK) model incorporating plasma, peripheral blood mononuclear cells, LN extracellular fluid, and LN intracellular compartments, and critically, a dynamic term accounting for changes in extracellular fluid (ECF) volume driven by lymphocyte trafficking. The model reveals that when fewer lymphocytes occupy the fixed LN volume, ECF volume expands, maintaining steeper drug concentration gradients that drive greater intracellular uptake. These findings identify lymphocyte trafficking as a previously unrecognized modulator of intranodal ARV distribution, one that is orthogonal to barrier-level penetration and operates through volume dynamics within the LN itself. Understanding these mechanisms is essential for optimizing ARV therapy in lymphoid reservoir sites and may inform strategies to enhance drug delivery to other pharmacological sanctuary sites.

## METHODS

### Animal model

CD-1 outbred mice (6-10 weeks old, Crl:CD1(ICR), Charles River) were used to optimize L-selectin functional antibody blocking (FAB) and ARV delivery. Mice were maintained according to Animal Use Protocols (AUP #1320 and #1367) approved by the Institutional Animal Care and Use Committee (IACUC) at the University of Delaware. Intravenous injections were performed following standard protocols. Full protocols for T lymphocyte isolation, CFSE labeling, and adoptive transfer are provided in Supplementary Methods. Briefly, primary T lymphocytes were isolated from mouse spleens via negative selection MACS and labeled with CFSE prior to intravenous transfer, as previously described^29^.

### Functional antibody blocking of L-selectin

To prevent lymphocyte ingress into the lobule, a functional antibody blocking (FAB) strategy using anti-mouse L-selectin (CD62L) antibodies (InVivoMAb, BioXCell) was employed via tail vein injection (**Figure S1**). Validation of functional lymphocyte blocking was conducted by testing doses from 0.004 to 8 mg/kg to identify a minimum functional FAB concentration. Using a dual-tail vein injection strategy, L-selectin antibody was injected into the left lateral tail vein, followed by 1.5x10^6^ CFSE-labeled T lymphocytes into the right lateral tail vein 2 hours later. Mice were monitored for 4 hours, euthanized, and blood was collected via cardiac puncture. Lymph nodes were collected, fixed, sectioned, and imaged using confocal microscopy. Dose-response data confirming effective FAB at 0.4 mg/kg are shown in **Figure S2**.

### ARV doses and sample preparation

EFV (38 mg/ml), RAL (70.1 mg/ml), and TDF (50 mg/ml) were prepared in sterile dimethylsulfoxide (DMSO). Samples prepared for injection used diluted stock solutions in a final 100 µL injection vehicle comprised of 30% DMSO and 70% PBS. This concentration of DMSO is safe for injection and allows for maximum solubility^30^. Samples were prepared by first warming stock solution, DMSO, and PBS to 60 ºC on a heat block. Solutions were mixed at appropriate volumes to get a desired final concentration in the 100 µL injection volume. After proper mixing of solution, samples were preloaded in a syringe, and allowed a slow ramp down to 37°C prior to injection. Doses were initially allometrically scaled from clinical doses according to a 75 kg human to appropriate mouse dose. From here, doses were scaled by factors from 2 to 10 times higher than this allometric dose due to limit of detection (LOD) in tissue fractions. Final administered doses were EFV = 24 mg/kg, RAL = 48 mg/kg, and TDF = 40 mg/kg, selected to maximize tissue detectability within established solubility and toxicity limits^31–34^.

### ARV injections with FAB

The ARV doses were consistently prepared without precipitate formation and were reliably detected in four pooled lymph nodes via mass spectrometry (MS). In each FAB experiment, five mice were used per ARV species: one untreated control, two mice for the control group (no FAB), and two mice for the experimental group (with FAB). Each of these experiments was conducted three times for each ARV species, involving a total of 3 untreated mice, 6 control mice, and 6 experimental mice. FAB experiments were performed identically for all ARV species (**Figure 1**). At t = 0, two of the mice received an L-selectin antibody injection at 0.4 mg/kg in the left lateral tail vein. After 2 hours, both experimental mice and two new control mice received an ARV injection into the right lateral tail vein, with doses listed in the **Supplementary Table S1**. After 4 more hours, the expected time constant of biodistribution, mice were euthanized, and blood and lymph nodes were collected.

**Figure 1:**
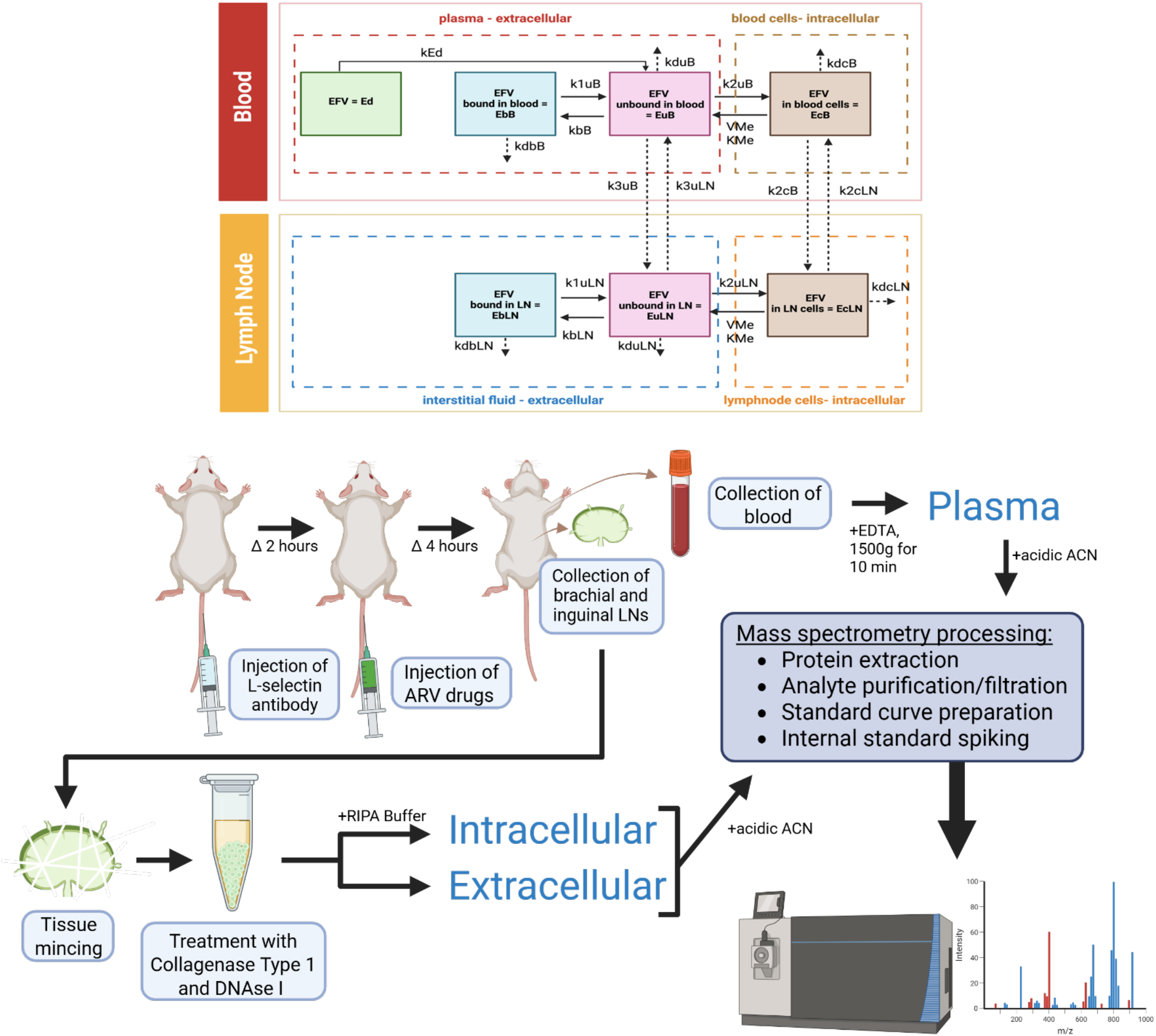
Schematic of experimental protocol for assessing cell-mediated ARV transport. Plasma, intracellular, and extracellular fractions are processed and analyzed via MS. (Biorender.com)

### Tissue collection and fractionation protocol

After euthanasia, blood was collected in a 0.1 M EDTA prefilled syringe, transferred to microcentrifuge tubes, and spun at 1500 x g for 10 minutes at 4°C to separate plasma from RBCs and leukocytes. Brachial and inguinal lymph nodes were pooled, submerged in 200 µL RPMI1640 with 5 mg/ml Collagenase Type I and 10 µg/ml DNAse I, and incubated for 1 hour at 37°C with periodic pipette agitation to dissociate the tissue. Cell counts were taken, and samples were centrifuged at 300 x g for 5 minutes to pellet the cells. Supernatants were collected as the extracellular (EC) fraction, and pellets were digested with 100 µL RIPA Buffer and 100 µL ddH2O to obtain the intracellular (IC) fraction. Samples were stored at -20°C until UPLC-MS/MS extraction.

### ARV extraction from tissue compartments

Tissue fraction samples were thawed, spun down, and a 150 µL subsample spiked with 2 µL ARV isotope and 148 µL cold acidic acetonitrile (ACN). After vortexing, centrifuging, and incubating at 50°C for 45 minutes, samples were filtered. An 80 µL aliquot was collected for mass spectrometry, with the remainder stored at -20°C. Mass spectrometry detection was performed using a Thermo Q-Exactive hybrid Quadrupole-Orbitrap mass spectrometer equipped with an UltiMate 3000 UHPLC system (Thermo Scientific), using heated electrospray ionization (HESI). Full instrumentation parameters are provided in **Supplementary Methods (Figure S4-S6)**.

### ARV extraction from plasma

Plasma samples were thawed, spun down, and a 150 µL subsample was spiked with 2 µL ARV isotope and 298 µL cold ACN. Following vortexing, centrifuging, and incubating, the supernatant was filtered. An 80 µL aliquot was collected for mass spectrometry, with the rest stored at -20°C.

### Tenofovir dephosphorylation

After cell uptake, TFV undergoes two phosphorylation steps to yield TFV diphosphate. To determine the entire TFV concentration in the intracellular compartment, an additional step was performed following extraction. Samples were dephosphorylated using potassium phosphate dibasic (Sigma) for 120 minutes at 37ºC, followed by chromatography as described in the literature^35^.

### Animal measurements and normalizations

To normalize data and correct for experimental variability, tissue sample measurements were collected for each mouse. Plasma MS concentrations were adjusted for the dilution factor of 100 µL based on the total blood volume drawn via intracardiac puncture. Cell counts from each intracellular (IC) fraction were multiplied by the approximate size of an isolated T cell (diameter ∼8 µm, volume ∼175 fL)^36^ to estimate intracellular lymph node volume, and IC MS concentrations were normalized to this volume. To account for any inconsistencies in intravenous injections, IC and extracellular (EC) fraction concentrations were normalized to plasma concentrations.

### Statistical analysis

All experiments were performed in triplicate for each ARV species (n = 6 mice per condition across 3 replicates). Results are presented as average ± standard deviation. Average concentrations between control and experimental groups were compared using a homoscedastic two-sample t-test, which assumes equal variance.

### Multi-compartment PBPK model

We developed nonlinear PBPK models by extending previously published PK models for the investigated drugs^21– 24^. The resulting PBPK models describe ARV distribution across four interconnected compartments: plasma (bound and unbound), peripheral blood mononuclear cells (PBMCs), LN extracellular fluid (bound and unbound), and LN intracellular (IC) space. The model incorporates mechanistic reaction kinetics while dynamically accounting for changes in ECF volume. ARVs in plasma can either diffuse into the LN or be taken up by lymphocytes and transported intracellularly. The model therefore captures both free-drug and cell-mediated transport and accounts for ECF volume changes as a concentration-modulating factor that influences drug distribution within the LN.

### Model assumptions

1. The protein binding and unbinding of ARV occur within the plasma and LN ECF compartments.
2. Protein binding and unbinding in the plasma is faster than in the LN ECF compartment.
3. The forward and backward drug transport rates from the blood plasma to the blood IC compartment are the same as those in the LN compartment.
4. The forward and backward drug transport rates from the IC blood to the IC LN compartment are the same.
5. The rate constant for drug transport from plasma to the LN ECF compartment was assumed to be the same as that from PBMC to the LN IC compartment.
6. Up to 50% of the total volume (*V*_*tot*_) in the LN can be replaced with ECF (interstitial fluid), denoted by *V*_*max*_.
7. ECF volume (*V*_*ex*_) is 20% of *V*_*tot*_ at steady state.

### Model structure by drug

Three drug-specific PBPK models were developed, each adapting a published human PK model^21–24^. Key structural features are summarized in **Table 1**, and model structures are illustrated in **Figure 2**. All three models share the ECF volume-dynamics term **(Eq. 1)**; drug-specific differences reflect the known PK properties of each compound.

**Table 1.** Summary of drug-specific PBPK model features.

| **Feature** | **Efavirenz** | **Tenofovir** | **Raltegravir** |
|---|---|---|---|
| Drug class | NNRTI | NRTI | INSTI |
| Dose | 600 mg once daily | 300 mg once daily | 400 mg twice daily |
| Route of administration | Oral | Oral | Oral |
| Published base PK model | Ref [21] | Ref [22, 23] | Ref [24] |
| Plasma protein binding | High (99.8%) | Minimal | Moderate (83%) |
| Plasma unbound fraction | 0.2% | High / mostly unbound | 17% |
| Bound/unbound plasma compartments | Yes | No | Yes |
| LN bound/unbound compartments | Yes | No | Yes |
| PBMC compartment | Yes | Yes | Yes |
| LN intracellular compartment | Yes | Yes | Yes |
| Nonlinear intracellular transport | Yes | No | No |
| Intracellular phosphorylation | No | Yes | No |
| Peripheral compartment retained | No | No | Yes |
| Dynamic LN ECF volume | Yes | Yes | Yes |
| Equations in Supplement | Eqs. S1–S7 | Eqs. S8–S16 | Eqs. S17–S24 |

**Figure 2.**
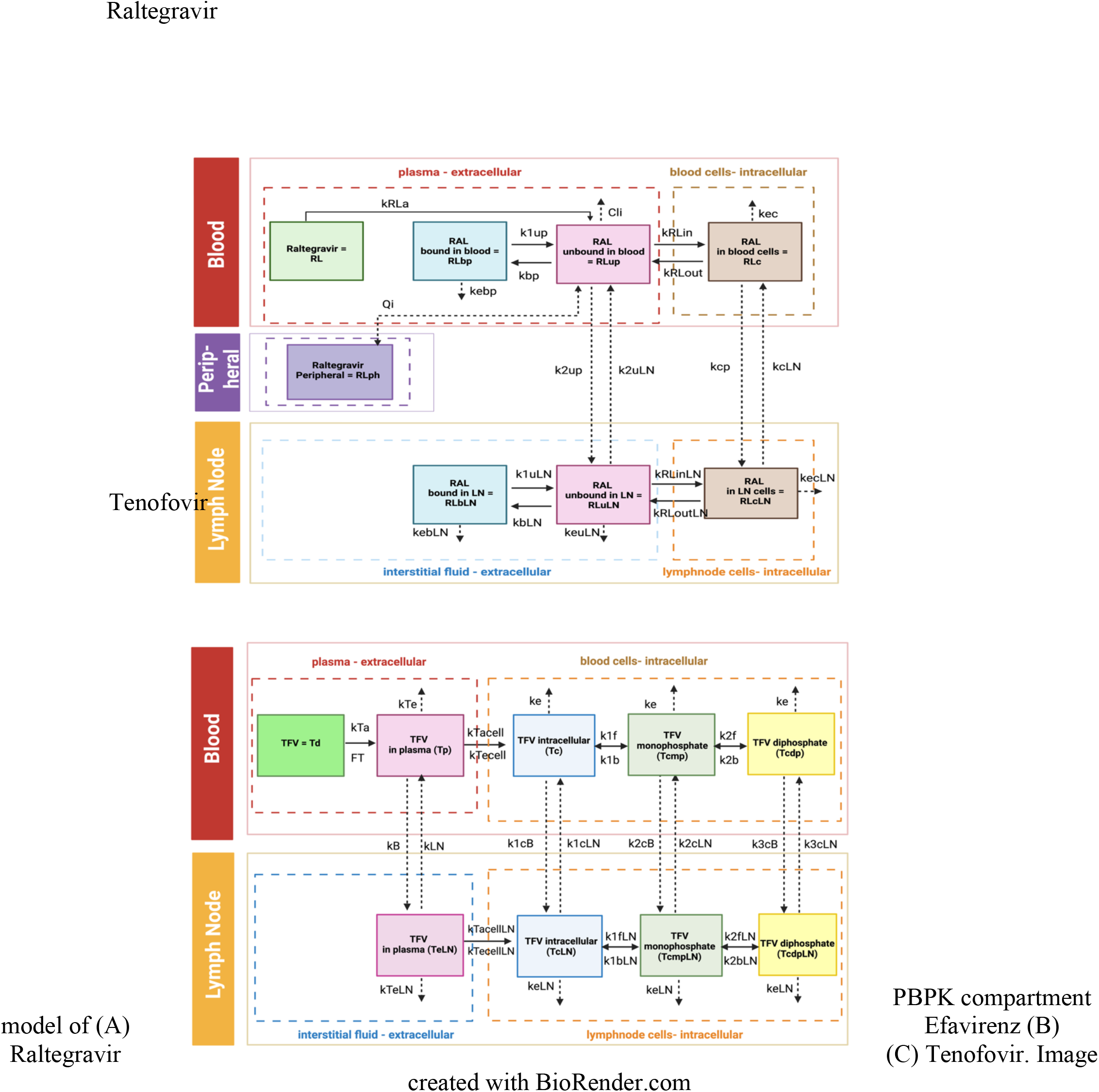
PBPK compartment model of (A) Efavirenz (B) Raltegravir (C) Tenofovir. Image created with BioRender.com

### The extracellular volume dynamics formulation

Key modeling contributions of this work include extending published human PK models into drug-specific PBPK frameworks, incorporating LN EC and IC drug compartments, representing drug-specific binding/unbinding kinetics, and coupling these models to dynamic LN ECF volume changes. Among these contributions, ECF volume dynamics within the LN is the central modeling contribution of this work because it allows lymphocyte trafficking–associated changes in LN volume to directly influence intranodal drug exposure. The equation below links lymphocyte trafficking to intranodal drug distribution and is applied uniformly across all three drug models:

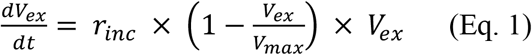

The EC volume equation (Eq. 1) is applied uniformly across all PBPK models presented in this study. To mimic the effect of FAB on lymphocyte entry into the LN, our model uses a logistic growth model (*V*_*ex*_) for the EC fraction of total volume in the LN. Logistic growth enforces a physiological upper bound where ECF cannot expand indefinitely within the fixed LN capsule, and *V*_*max*_ (50% of total LN volume) captures this constraint. By tuning the ECF volume (*V*_*ex*_) via the intrinsic growth rate (*r*_*inc*_), we controlled the modeled effect of lymphocyte entry into the LN. When *r*_*inc*_ = 0, *V*_*ex*_ remains at baseline (20% of LN volume), corresponding to normal lymphocyte trafficking. When *r*_*inc*_ = 3, *V*_*ex*_ expands toward *V*_*max*_, modeling the reduced cellular occupancy under FAB conditions. The time-varying *V*_*ex*_ term was then incorporated into the LN compartment equations (**Supplementary Eqs. S5–S7, S13–S16, and S22–S24**), allowing changes in LN ECF volume to influence intranodal drug concentrations through volume scaling and dilution effects.

### Model parameterization

The models were parameterized using published human PK data^21–24^. Sensitivity analysis was performed to evaluate the influence of key parameters on model predictions, with results shown in **Figures S10–S12**. All parameter values are provided in **Tables S2–S4**. Direct fitting to the murine experimental data was not performed; instead, the model was used to determine whether the ECF volume dynamics mechanism (Eq. 1) qualitatively reproduces the direction of the experimental observations. Quantitative species-specific validation is an acknowledged limitation and is discussed in the Discussion.

## RESULTS

Blocking lymphocyte entry into the LN paradoxically increased IC ARV concentration for two of three drugs tested. For each drug, we present the experimental observation, the apparent paradox, and the model-derived mechanistic explanation.

### FAB validation

To determine the viability of adoptive T lymphocyte transfer, 1.5x10^6^ CFSE-labeled T lymphocytes were intravenously injected into mice (**Figure 3A-B**). After 2 hours, mice were euthanized and brachial, inguinal, and popliteal lymph nodes were collected^37^. Tissues were then sectioned and imaged via confocal microscopy (**Figure 3C-D**). In all three lymph nodes representing three different locations in the body, strong labeled lymphocyte uptake was seen (**Figure S2A-D**). When counterstained with phalloidin, labeled T lymphocytes were shown to interact with HEVs in the paracortex region and were found in the avascular spaces within the lobule (**Figure S2E**). Immunostaining with anti-CD3 antibody confirmed that the FAB strategy did not cause issues with lymph node architecture or hypoplasia (**Figure S2F**).

**Figure 3:**
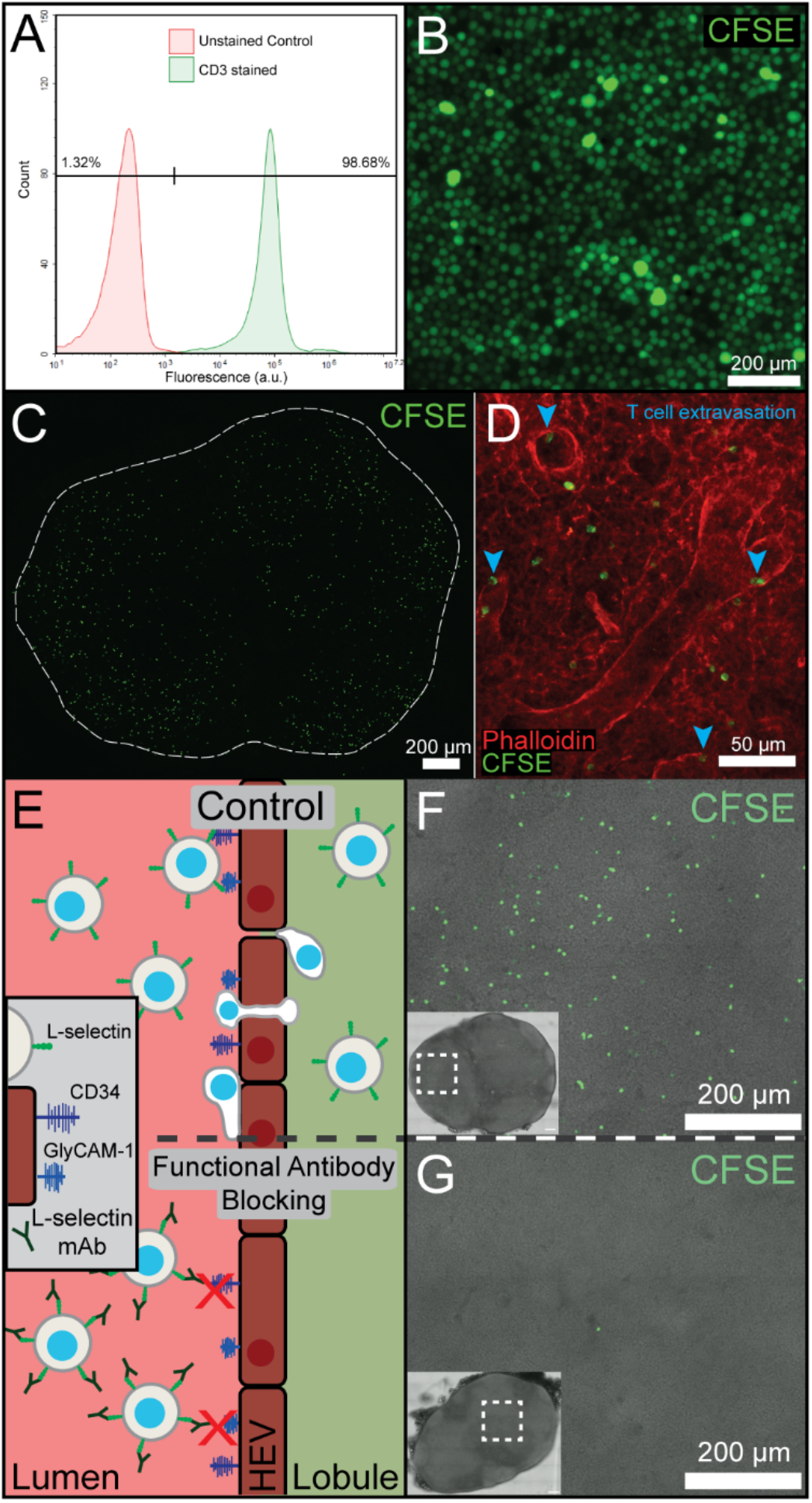
A) Validation of T lymphocyte isolation via flow cytometry. B) CFSE ex vivo labeling of T lymphocytes and C) adoptive cell transfer. Imaging of LN cross section and D) phalloidin staining confirm cell extravasation into LN lobule. E) Illustrative schematic and fluorescent imaging of F) control and G) anti L-selectin functional antibody blocking (FAB).

For the optimization of our L-selectin FAB scheme, a range of antibody doses were assessed (0.004-8 mg/kg). To confirm injection quality, adoptively transferred T lymphocytes were still detectable in blood at 2 hours. In tissue sections, it was clear that all doses below 0.4 mg/kg resulted in strong lymph node homing similar to the control animal **(Figure S3)**. However, at doses of 0.4 mg/kg and greater, lymphocyte ingress was greatly reduced, and less than one labeled T lymphocyte was identified per ROI. For further FAB experiments, the 0.4 mg/kg dose was selected.

### ARV extraction and detection

An optimized extraction and purification method was used to detect ARV species concentrations within the plasma and intracellular (IC) and extracellular fractions (EC). First, conventional protein precipitation techniques using ice-cold acidic acetonitrile were performed on all collected fractions. However, due to the logP and protein-bound fractions (PBF) of a number of the drugs, standard methods were insufficient. A heat shock step was added to the processing pipeline to denature proteins bound to ARV species. Using standard methods, EFV and RAL had recovery percentages of 9.9% and 15.5%, respectively. After incorporation of heat shock methodology, extraction efficiency increased for EFV and RAL to 96% and 94%, respectively. The stability of drugs through the heat treatment was confirmed experimentally.

### Tenofovir: FAB increases intracellular LN concentration by 88%

TFV was detected via Orbitrap MS using negative ion mode with a retention time of 2.15 minutes. Prior to submission, TFV IC compartments were dephosphorylated, and plasma and EC compartments were handled similarly to the other ARV species. After tissue normalizations (Methods), the plasma fraction, IC fraction, and EC fraction were calculated for control mice and FAB mice (**Table 2, Figure S7**). Plasma concentrations were equivalent between groups (10.362 vs. 10.547 µM; p = 0.9157), confirming that the IC difference reflects intranodal redistribution rather than differences in systemic drug delivery. If blocking lymphocyte entry reduces cell-mediated ARV transport into the LN, intracellular concentration should decrease, not increase. The 88% increase (p<0.05) in normalized IC TFV (9.063 → 17.047 µM) under FAB is therefore paradoxical and requires a mechanistic explanation (**Figure 4A**).

**Table 2.** Summary of measured ARV concentrations in tissue (average ± standard deviation).

|  |  | **Control** |  |  | **FAB** |  |  |
|---|---|---|---|---|---|---|---|
| **Drug Species** | **Sample Type** | **Plasma (µM)** | **IC (µM)** | **EC (µM)** | **Plasma (µM)** | **IC (µM)** | **EC (µM)** |
| Tenofovir | Raw Tissue | 5.528 (±1.665) | 391.96 (±255.789) | 788.712 (±32.941) | 5.730 (±1.755) | 622.25 (±161.654) | 755.228 (±17.675) |
| Efavirenz |  | 4.272 (±0.956) | 247.472 (±60.001) | 156.962 (±97.830) | 3.517 (±1.146) | 294.182 (±120.866) | 68.992 (±41.449) |
| Raltegravir |  | 4.126 (±0.676) | 1892.92 (±158.095) | 6100.143 (±345.442) | 4.019 (±0.644) | 1725.305 (±228.345) | 5928.398 (±647.051) |
| Tenofovir | Tissue Normalization | 10.362 (±2.144) | 9.063 (±5.04) | 0.7887 (±0.0329) | 10.547 (±3.565) | 17.047 (±4.219) | 0.755 (±0.0177) |
| Efavirenz |  | 8.263 (±1.148) | 6.967 (±1.874) | 0.1570 (±0.0978) | 5.768 (±1.547) | 6.156 (±2.507) | 0.0816 (±0.0309) |
| Raltegravir |  | 6.956 (±1.503) | 53.121 (±6.095) | 6.100 (±0.345) | 7.159 (±1.614) | 51.923 (±11.554) | 5.928 (±0.647) |
| Tenofovir | Plasma Normalization |  | 0.841 (±0.441) | 0.0762 (±0.0182) |  | 1.755 (±0.617) | 0.0832 (±0.0451) |
| Efavirenz |  |  | 0.847 (±0.223) | 0.0185 (±0.00962) |  | 1.063 (±0.325) | 0.0140 (±0.00361) |
| Raltegravir |  |  | 8.043 (±2.336) | 0.928 (±0.291) |  | 7.564 (±2.670) | 0.853 (±0.151) |

**Figure 4:**
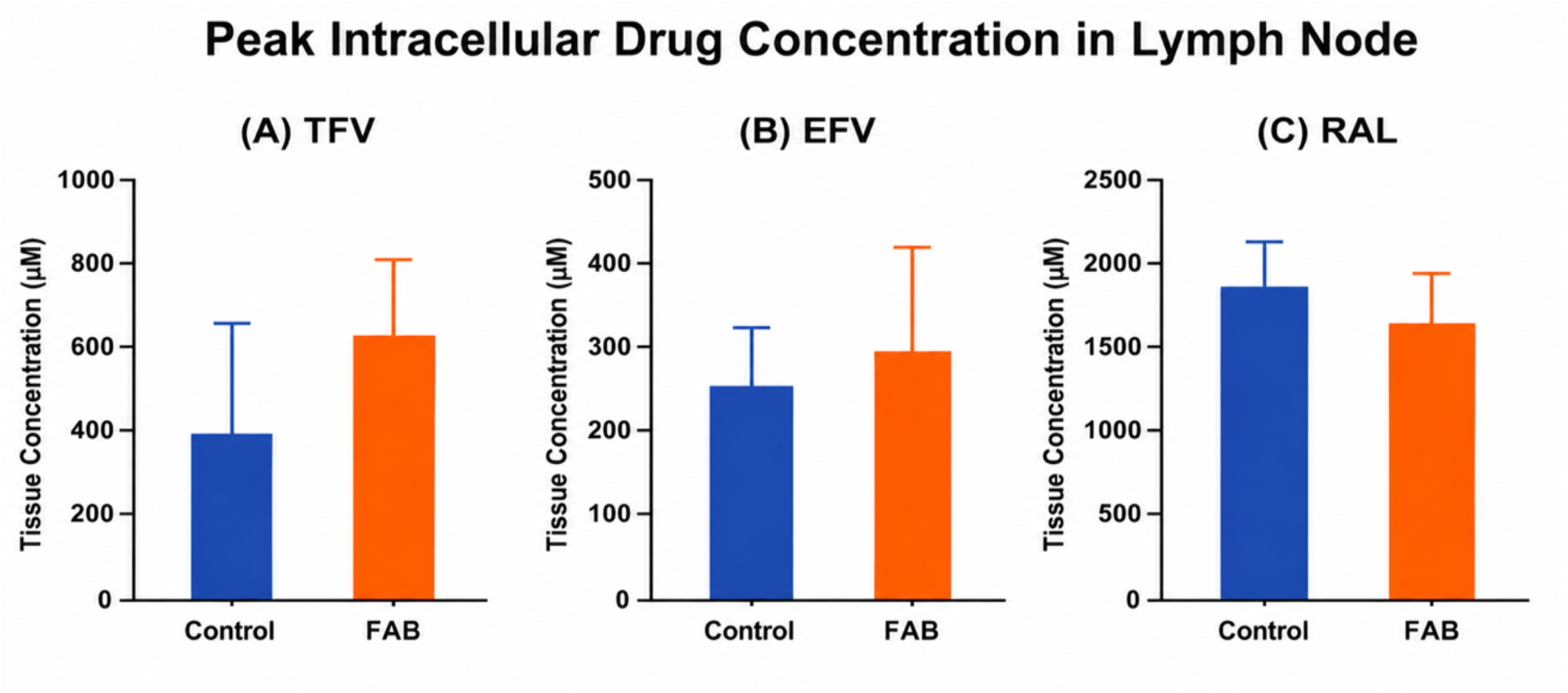
Murine experimental results showing intracellular lymph node concentrations of (A) Tenofovir (TFV) (B) Efavirenz (EFV), and (C) Raltegravir (RAL) under control and FAB conditions.

The computational model resolves this paradox through ECF volume dynamics. When lymphocyte entry is blocked (*r*_*inc*_ = 3), *V*_*ex*_ expands from its 20% baseline toward *V*_*max*_ (50% of LN volume). The larger extracellular compartment sustains higher ECF drug concentrations, steepening the ECF-to-IC gradient and driving greater IC uptake. Simulation predicts an increase in IC TFV concentration under FAB conditions, consistent with the experimental observation **(Figure 5A, Figure S13A)**. The model further predicts that the net total LN TFV concentration slightly decreases under FAB conditions, because the rise in IC drug is outweighed by the dilution of ECF drug across the expanded extracellular space (Figure S13B and C). This prediction is consistent with the near-unchanged EC fraction (0.7887 vs. 0.755 µM). TFV’s irreversible IC phosphorylation to TFV-DP amplifies this gradient-driven uptake, making it the most sensitive of the three drugs to ECF volume dynamics.

**Figure 5:**
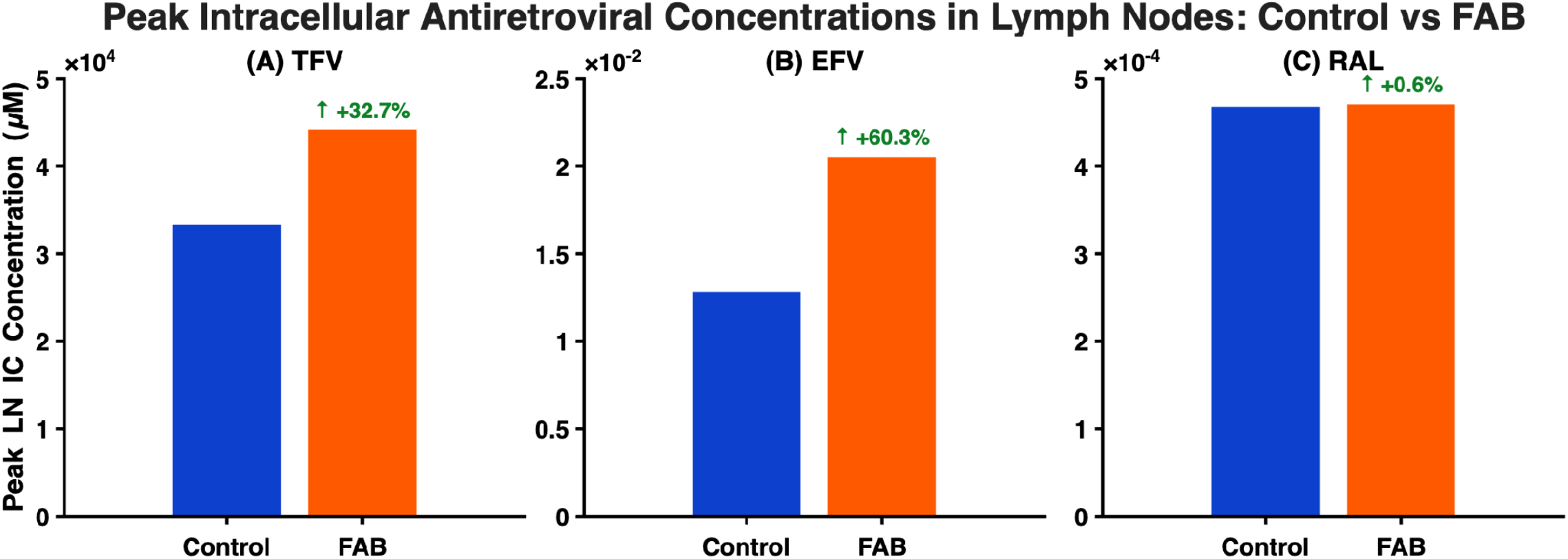
Human data simulation results showing peak IC LN concentrations of (A) Tenofovir (TFV), (B) Efavirenz (EFV), and (C) Raltegravir (RAL) under control and FAB conditions.

### Efavirenz: FAB increases plasma-normalized LN concentration; raw IC data require interpretation

EFV was detected via Orbitrap MS using negative ion mode with a retention time of 3.33 minutes. After tissue normalizations, the plasma fraction, IC fraction, and EC fraction were calculated for control mice and FAB mice (**Table 2, Figure S8**). Normalized IC EFV concentrations were numerically lower in FAB mice (6.156 vs. 6.967 µM), while plasma concentrations were also lower in the FAB group (5.768 vs. 8.263 µM). The plasma discrepancy was unexpected, whereby the model predicts no significant difference in plasma EFV between groups and may reflect injection variability or inter-animal PK differences unrelated to FAB. The plasma-normalized IC:EC ratio (75.9 vs. 45.8 for FAB vs. control) is consistent with the TFV finding, with relative IC accumulation increasing when lymphocyte entry is blocked (**Figure 4B**).

The same ECF volume expansion mechanism explains the EFV finding. Under FAB condition, simulation predicts an increase in LN IC EFV concentration **(Figure 5B, Figure S13D)**. The model also predicts that LN ECF EFV decreases, whereas total LN EFV remains nearly unchanged (Figure S13E and F). The increase in LN IC EFV under FAB was largely offset by the reduction in the ECF compartment, resulting in only a negligible net change in total LN EFV concentration. Thus, FAB did not increase overall EFV accumulation in the LN; instead, it shifted the relative distribution toward the IC space, similarly to TFV and consistent with the experimentally observed increase in the IC:EC ratio. However, we expect that EFV’s high lipophilicity and reversible membrane partitioning facilitate exchange between LN ECF and IC spaces, which is captured by the protein-binding and bidirectional transport terms in the EFV model, and support the FAB-induced shift toward the IC compartment.

### Raltegravir: FAB produces no significant change in intracellular concentration

RAL was detected via Orbitrap MS using positive ion mode with a retention time of 2.62 minutes. After tissue normalizations, the plasma fraction, IC fraction, and EC fraction were calculated for control mice and FAB mice (**Table 2, Figure S9**). IC concentrations were statistically indistinguishable between groups, as were the normalized ratios (**Figure 4C**). The absence of a detectable FAB effect on IC RAL is consistent with relatively rapid, reversible partitioning between EC and IC compartments and limited IC retention. IC RAL concentrations closely track plasma concentrations over time while remaining substantially lower than plasma concentrations, suggesting limited IC accumulation. These kinetic characteristics suggest that RAL is less prone to sustained IC accumulation than drugs with greater IC retention.

This interpretation is supported directly by the model. Simulated RAL IC concentrations remained nearly unchanged between control and FAB conditions, despite modest transient differences in the ECF and total RAL concentrations (**Figure 5C, Figure S13G-I**). The Raltegravir IC null result thus constrains the ECF volume-dynamics mechanism, indicating that the effect of blocking lymphocyte entry is not universal but depends on drug-specific transport and retention kinetics across compartments. Drugs that equilibrate slowly (irreversible phosphorylation, as in TFV; high protein binding, as in EFV at 99.8%) are sensitive to ECF volume shifts; drugs that equilibrate rapidly (Raltegravir) are not.

## DISCUSSION

The high rate of lymphocyte trafficking into and out of LNs, coupled with known rates of ARVs drug uptake, suggests that lymphocytes may act as incidental carriers of ARV drugs. To test this hypothesis, we performed murine experiments in which lymphocytes’ entry into the LN was selectively prevented using FAB scheme. Unexpectedly, we observed that IC drug concentration within the LN increased substantially under conditions of blocked lymphocyte entry for TFV and EFV. To further investigate this paradoxical finding, we developed computational PBPK models parameterized with clinical pharmacokinetic data. Both experimental and simulated results showed consistent patterns: blockade of lymphocyte entry led to higher IC drug accumulation within the LN for both compounds.

We hypothesized that this higher IC drug concentration paradox can be explained by considering the balance between lymphocytes, ECF, and drug availability within the fixed volume of the LN. The intuition is as follows: imagine the LN as a bucket, where the bucket’s contents consist of lymphocytes (cells) and ECF. The total bucket volume is constant, so by pressure homeostasis, a decrease in one component necessarily leads to an increase in the other. When lymphocyte entry is blocked, some of the space normally occupied by cells fills with ECF (**Figure 6**). The enlarged ECF compartment sustains higher drug concentrations in the interstitial space, steepening the ECF to IC concentration gradient and driving increased IC uptake over the 4-hour observation window. This sustained exposure enables lymphocytes to accumulate more drug, thereby increasing the IC drug concentration across the LN. Our computational model resolved this by explicitly incorporating extracellular volume dynamics (Eq. 1) through the influx parameter *r*_*inc*_, which is used as a switch to represent lymphocyte entry conditions. When *r*_*inc*_=0 (normal trafficking), EC volume (*V*_*ex*_) remained at its 20% normal baseline; when *r*_*inc*_= 3 (FAB conditions), *V*_*ex*_ expands logistically toward *V*_*max*_ (50% of LN volume), sustaining the gradient that drives IC drug accumulation. This mechanism operates within the LN itself, downstream of drug entry across the HEV barrier, and is therefore orthogonal to the barrier-level transport failures that have been the primary focus of prior modeling work.

**Figure 6:**
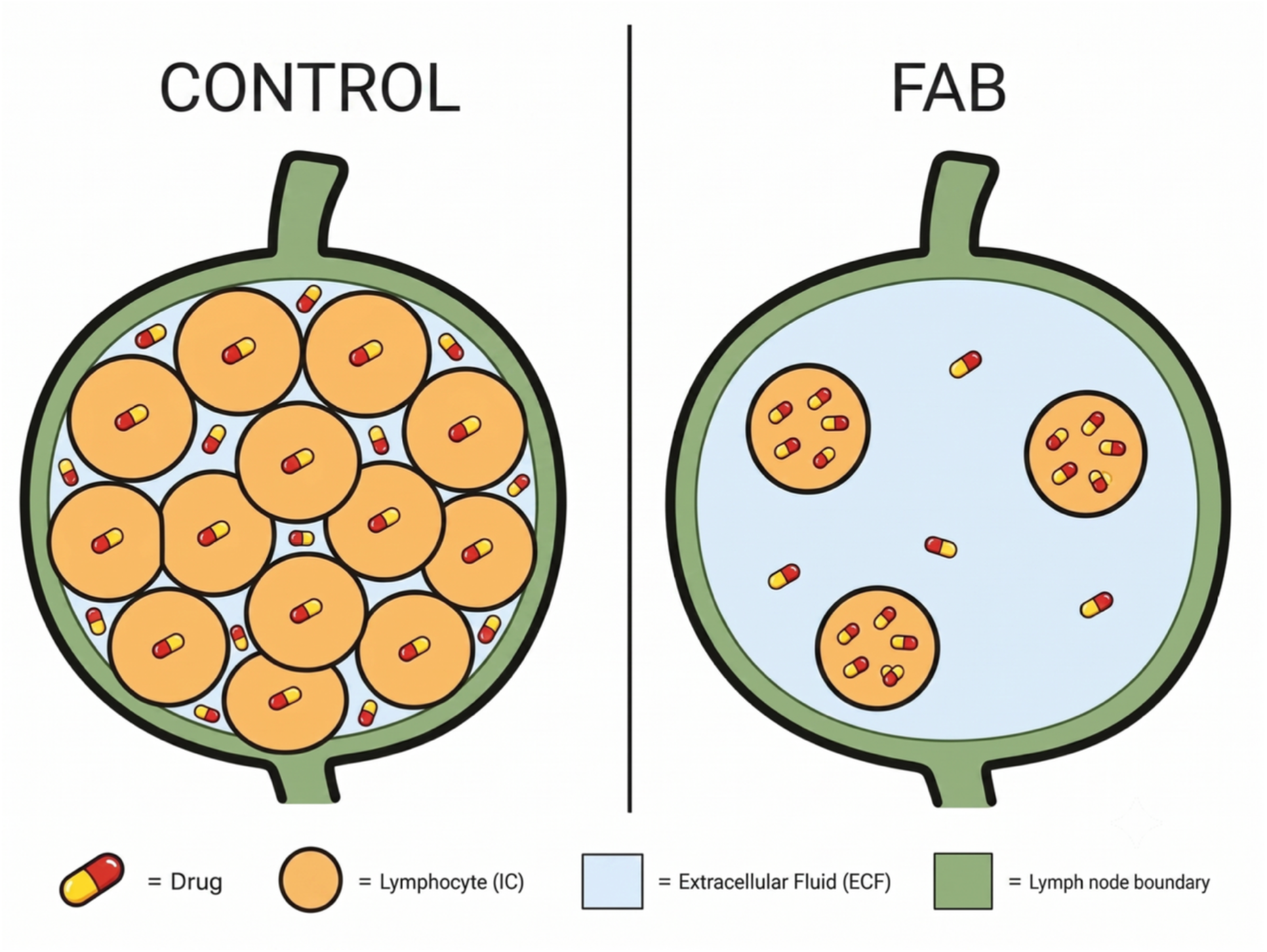
Pictorial representation of how blocking the lymphocyte population impacts the extracellular fluid (ECF) and intracellular (IC) compartments of the LN and drug distribution. Image created via Gemini.

This work extends a body of PK modeling that has treated LN drug penetration as a fixed property of the barrier. Sanche et al. (2017) demonstrated computationally that static penetration failure across the HEV is sufficient to drive virological failure and resistance emergence within the LN. Their model correctly identifies the HEV barrier as the primary obstacle to LN drug access but treats intranodal drug distribution as uniform once the drug enters the compartment. The ECF volume dynamics mechanism identified here operates at a different level: it governs how the drug distributes within the LN once it has crossed the barrier. Together, these two mechanisms of barrier-level penetration and intranodal ECF volume dynamics provide a more complete picture of why LN drug concentrations are both low and variable across patients. The present findings are consistent with Sanche et al. (2017) in clinical implications, whereby limited LN drug access degrades therapeutic efficacy but identifies an additional, modifiable determinant of intranodal distribution^17^.

The null result for RAL constrains the ECF volume dynamics mechanism and provides a mechanistic probe of its drug-class dependence. RAL IC concentrations were statistically indistinguishable between FAB and control mice, and the model likewise predicts negligible changes in RAL IC exposure under FAB conditions. This insensitivity is consistent with RAL’s rapid equilibration between ECF and IC compartments. Because RAL redistributes quickly in response to ECF concentration changes, any gradient advantage created by ECF expansion is rapidly dissipated. Sensitivity to ECF volume shifts is therefore governed by the reversibility of IC drug retention, which is a drug-class-level property. By contrast, TFV undergoes irreversible IC sequential phosphorylation to tenofovir monophosphate and then to tenofovir diphosphate, which traps the drug intracellularly and allows the sustained ECF-to-IC gradient to accumulate over time. EFV’ s high protein binding (99.8%) similarly slows equilibration. The RAL, TFV, and EFV FAB results are quantitative predictions of the models, which were confirmed experimentally. Drugs with slow IC equilibration kinetics are sensitive to ECF volume shifts; drugs with rapid equilibration are not.

There are two important limitations of this study that warrant explicit acknowledgment. First, all experiments were conducted at a single time point (4 hours post-ARV injection), corresponding to the expected biodistribution time constant. Time-course data would strengthen the mechanistic interpretation by confirming that IC concentrations rise progressively under FAB conditions. Secondly, the computational model was parameterized using published human PK data; direct fitting to the murine experimental data was not performed. The model is used here to identify the qualitative direction of the mechanism, not to make quantitative species-specific predictions.

The clinical relevance of these findings lies in the complexity of achieving and maintaining optimal ARV concentrations within lymphoid tissues. Although complete prevention of lymphocyte trafficking may not commonly occur in clinical practice, impaired lymphocyte entry may represent an important limiting condition for understanding how cell-mediated transport contributes to ARV distribution within lymph nodes. By reducing lymphocyte trafficking, this study provides insight into how changes in cellular occupancy can alter IC ARV concentrations within lymphoid compartments. These findings further suggest that intranodal drug distribution may depend on the interplay between extracellular drug transport, intracellular partitioning, and lymphocyte-associated transport rather than on passive diffusion alone. Changes in the relative distribution of drug between extracellular and intracellular compartments may therefore influence local pharmacologic exposure within the LN. While this work did not directly examine inflammation-driven increases in lymphocyte trafficking, this remains an important area for future investigation. Future studies could extend this framework to determine whether elevated lymphocyte influx alters ARV penetration within LN lobules and contributes to heterogeneous drug exposure in lymphoid tissues.

## CONCLUSION

This experimental and mathematical study underscores the critical role of lymphocyte trafficking in modulating the IC concentration of ARVs within the LN. Our findings show that blocking lymphocyte entry increases IC drug concentration for EFV and TFV, with no significant effect on RAL, a drug-class difference consistent with model predictions based on IC equilibration kinetics. The mechanistic explanation lies not in lymphocyte-mediated drug transport, but in ECF volume dynamics that result from changes in LN cellular occupancy. When fewer lymphocytes occupy the fixed LN volume, ECF expands, sustained drug gradients drive greater IC uptake, and IC concentrations rise. These results suggest that modulating lymphocyte trafficking can affect ARV exposure within LN tissues. Therefore, integrating lymphocyte trafficking dynamics into ART design could lead to more effective strategies for targeting HIV reservoirs in lymphatic tissues.

## SUPPORTING INFORMATION

Supplemental materials can be found in the accompanying file _*“*_Supplemental Material_FAB Paper.pdf_*”*_

## ACKNOWLEDGEMENTS

Manuscript figures were generated using BioRender and Gemini.

## DATA AVAILABILITY STATEMENT

Data and materials supporting this study’s findings are available from the corresponding author upon reasonable request.

