## Supplemental Material for "Effects Of Lymphocyte Trafficking on Antiretroviral Drug Distribution within the Lymph Node"

**EXPERIMENTAL SUPPLEMENTAL METHODS AND FIGURES**

#### METHODS

##### Reagents and preparation of ARV stocks

The following chemicals were provided by Merck: Efavirenz (EFV) and Raltegravir (RALT). Each drug was dissolved in dimethylsulfoxide (DMSO). A stock of EFV (38 mg/mL) and RALT (70 mg/mL) were prepared and filtered through a ultrafiltration microcentrifuge tube (3kDA MWCO or 0.22 µm, CellTreat) at 10,000 rcf for 2 minutes. Stocks were aliquoted and stored in -80°C until use.

ARV isotopes were acquired from the following manufacturers: AlsaChim (Illkirch-Graddenstaden, France); [$^{13}$C6]-Efavirenz ($^{13}$C6-EFV) and Raltegravir-d4 (RALT-d4). A 1 mg/mL stock solution of each ARV isotope was made in HPLC grade methanol.

For all ARV drugs a calibration curve of 100, 200, 400, 600, 800, and 1000 nM was submitted with each experimental run. Calibration curves were freshly prepared and submitted with each experimental run. Similarly, internal standards of ARV isotopes were prepared and added to all samples submitted, with the exception of ISL.

##### *In vivo* ARV delivery to murine model

CD1 mice (Charles River) ages 6-8 weeks were used as our murine model. For all ARV experiments, five mice were used per run: one untreated mouse, two mice for control group and two mice for functional antibody blocked group. Each ARV was tested thrice. Samples were prepared under sterile conditions. For each drug, a dose of, 24.8 mg/kg EFV, 48 mg/kg RALT, or 40 mg/kg ISL was delivered to each mouse.

Supplementary Table S1

| Antiretroviral Species | Drug Class | logP | PBF | Allometric scaling | Dose (mg/kg) |
|---|---|---|---|---|---|
| Efavirenz | Non-nucleoside RT Inhibitor | 4 | 0.99 | 3X | 24 |
| Raltegravir | Integrase Inhibitor | 1.1 | 0.8 | 6X | 48 |
| Tenofovir DF | Nucleotide RT Inhibitor | -1.6 | 0.01 | 10X | 40 |

##### Functional antibody blocking of L-selectin in murine models

To ensure that changes in drug concentration between control groups and obstructed lymphocyte groups are cell mediated, a range of antibody concentrations of anti-L-selectin monoclonal functional antibody blocking (CD62L, BioxCell) was tested and determined for optimal dosing. A dose of 0.4 mg/kg was selected based on these preliminary studies.

Briefly, each mouse was weighed to prepare a L-selectin functional antibody blocking treatment at 0.4 mg/kg. Anti-L-selectin samples were prepared in 1 mL slip tip syringes (BD) with a 27 G needle. Mice were immobilized in an animal restraint and placed under a heat lamp and warmed until tail vein was visible (30-60 seconds). Tails were first cleaned with alcohol wipes and then L-selectin antibody blocking samples were delivered via tail vein injection and mice were left for four hours before ARV delivery.

ARV doses were determined by using approximate human doses and scaling down to reflect mice weight. Control mice were weighed, and ARV doses were calculated and prepared from stock ARV solutions diluted to100 μL in 1X phosphate buffered saline (PBS), per institutional animal care and use committee (IACUC) regulations. Two hours after functional antibody blocking treatment of the blocked mice group, mice were again restrained, and ARV doses were delivered to control and blocked mice via tail vein injection.

**Lymph node collection**

Four hours after ARV treatment, mice were euthanized in $CO_2$ chambers, followed by cervical dislocation. Due to the size of the tissue, poor drug penetration, and limit of detection by MS, four lymph nodes (two inguinal and two brachial) from each mouse were harvested and pooled for analysis.

**Fractionation of intracellular and extracellular tissue compartments**

To isolate the extracellular compartment, the lymph nodes were transferred to 1.5 mL microcentrifuge tubes, minced, and treated with 200 μL of a extracellular matrix (ECM) digestion solution containing 5 mg/mL collagenase type 1 (Worthington) and 10 μg/mL DNAse 1 (Worthington) for 1 hour at 37 °C. Digested samples were centrifuged at 300 rcf for 5 minutes at 4°C. The supernatant was collected as our extracellular compartment. The remaining cell pellet is then lysed in 200 μL of Pierce RIPA buffer (Thermo) for 10 minutes and saved as our intracellular compartment. Samples were stored in -20°C until ARV extraction.

**Plasma collection**

Blood was collected from each sacrificed mouse via cardiac puncture. Syringes were prepped with 100 μL of 0.1M aqueous ethylenediaminetetraacetic acid (EDTA, Sigma) to prevent coagulation of blood. The blood samples were transferred to 1.5 mL microcentrifuge tubes and centrifuged at 1500 rcf for 10 minutes at 4°C to sediment blood cells. The supernatant was collected and saved as our plasma fraction. Samples were stored in -20°C until ARV extraction.

**ARV extraction from tissue compartments**

A 150 μL aliquot of each tissue fraction was transferred to a new 1.5 mL centrifuge tube. Then samples were dosed with 2 μL of the respective ARV isotope as an internal standard and treated with 148 μL of acidic acetonitrile. Samples were vortexed thoroughly for 15 seconds and then centrifuged at 1500 rcf for 1 minute. Samples were then placed on a Thriller thermoshaker incubator block (Peqlab) set at 50°C and mixed at 350 rpm for 45 minutes to disassociate protein bound drug species. Samples were allowed to slightly cool before they were vortexed thoroughly followed by a quick spin. Samples were transferred to a 500 µL Spin-X ultrafiltration microcentrifuge tube (10 kDa MWCO, Corning) and centrifuged at 4,500 rcf for 10 minutes. An 80 µL aliquot of each filtrate was collected and submitted for mass spectrometry. The remaining samples were preserved and stored at -20°C.

**ARV extraction from plasma fraction**

Similar to tissue processing, a 150 µL aliquot of each plasma sample was transferred to a new 1.5 mL centrifuge tube. Samples were dosed with 2 µL of the respective ARV isotope as an internal standard and treated with 298 µL of acidic acetonitrile (0.1 M formic acid in ultrapure water). Samples were vortexed thoroughly for 15 seconds and then centrifuged at 1500 rcf for 1 minute. Samples were then placed on a Thriller thermoshaker incubator block (Peqlab) set at 50°C and mixed at 350 rpm for 45 minutes to disassociate protein bound drug species. Samples were allowed to slightly cool before they were vortexed thoroughly followed by a quick spin. Samples were transferred to a 500 µL Spin-X ultrafiltration microcentrifuge tube (10 kDa MWCO, Corning) and centrifuged at 4500 rcf for 10 minutes. An 80 µL aliquot of each filtrate was collected and submitted for mass spectrometry. The remaining samples were preserved and stored at -20°C.

**Tenofovir dephosphorylation**

After being taken up by cells, tenofovir undergoes through two phosphorylation steps to yield tenofovir diphosphate. In order to determine the entire tenofovir concentration in the intracellular compartment, an additional step was performed following extraction. Samples were dephosphorylated using potassium phosphate bibasic (Sigma) for 120 minutes at 37°C followed by chromatography as described in literature.

**Instrumentation**

Mass spectrometry (MS) detection was performed using a Thermo Q-Exactive hybrid Quadrupole-Orbitrap mass spectrometer (Thermo Scientific, Bremen, Germany) equipped with an UltiMate 3000 ultra high-performance liquid chromatography (UHPLC) system (Thermo Scientific, Germering, Germany), using conventional heated electrospray ionization (HESI).

**UPLC-MS/MS**

The thermostatted autosampler was maintained at 5°C with the column compartment temperature set at 40°C. Each sample had an injection volume of 10 µL and eluted through a ACQUITY UPLC CSH C18 (130 Å, 1.7µm, 2.1 mm x 30 mm) with an ACQUITY UPLC CSH C18 guard column (130 Å, 1.7µm, 2.1 mm x 5 mm) using a mobile phase gradient with a run time of 6 minutes. Mobile phase A: 0.1% (v/v) formic acid in LC-MS grade water; mobile phase B: 0.1% (v/v) formic acid in LC-MS grade acetonitrile. At a flow rate of 0.5 mL/min, the mobile phase gradient began at 95%:5% A:B, ramping to 5%:95% A:B from 0 to 4 minutes, remining at 5%:95% from 4 to 5 minutes, then returned to 95%:5% at 5.1 minutes, and remained until 6 minutes.

The following parameters were used for positive HESI mode: capillary spray voltage was set at 3.75 kV; sheath gas flow set at 45 psi, auxiliary gas flow at 10 arbitrary units (AU), sweep gas flow at 2 AU; the capillary temperature at 250°C, S-lens radiofrequency level to 50 rf, and auxiliary gas heater to 400°C. The following parameters were used for negative HESI mode: capillary spray voltage was set at -3.5 kV; sheath gas flow set at 45 psi, auxiliary gas flow at 10 AU, sweep gas flow at 2 AU; the capillary temperature at 275°C, S-lens radiofrequency level to 50 rf, and auxiliary gas heater to 400°C.

Efavirenz was detected in negative mode; all the other ARVs in this study (Raltegravir and Tenofovir) were detected in positive mode.

Analytes were monitored using targeted selected ion monitoring, data-dependent MS/MS mode (t-SIM-ddMS2). The following SIM parameters were used: resolution set to 17, 500, automatic gain control (AGC) target set at $5x10^4$, maximum injection time (IT) set to 100 ms, and isolation window 4 *m/z*. The following ddMS2 parameters were used: resolution set to 17, 500, AGC target at $2x10^5$, maximum IT to 100 ms, isolation window 4 *m/z*, and higher-energy collisional dissociation at normalized collision energy of 35 eV. Methodology for each drug was developed by Dr. Papa Nii Asare-Okai. Samples were run by mass spectrometry core personnel. Xcalibur software (Thermo Scientific) was used for data acquisition and analysis.

#### RESULTS

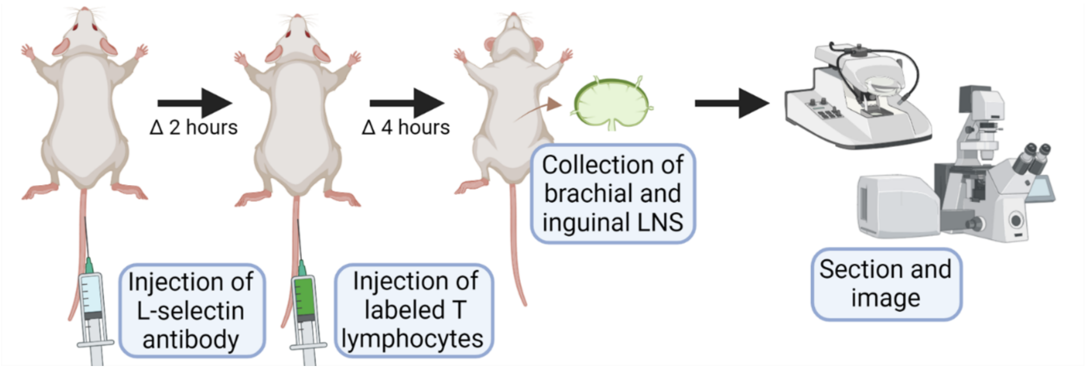


**Figure S1:** A cartoon overview of functional antibody blocking process.

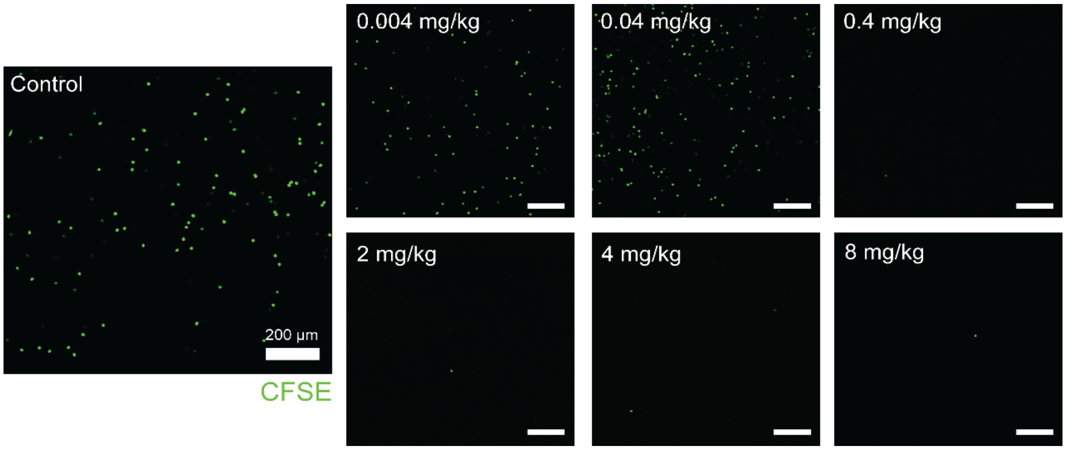


**Figure S2:** Adoptive T lymphocyte transfer following FAB shows doses at and above 0.4 mg/kg sufficiently block lymphocyte ingress.

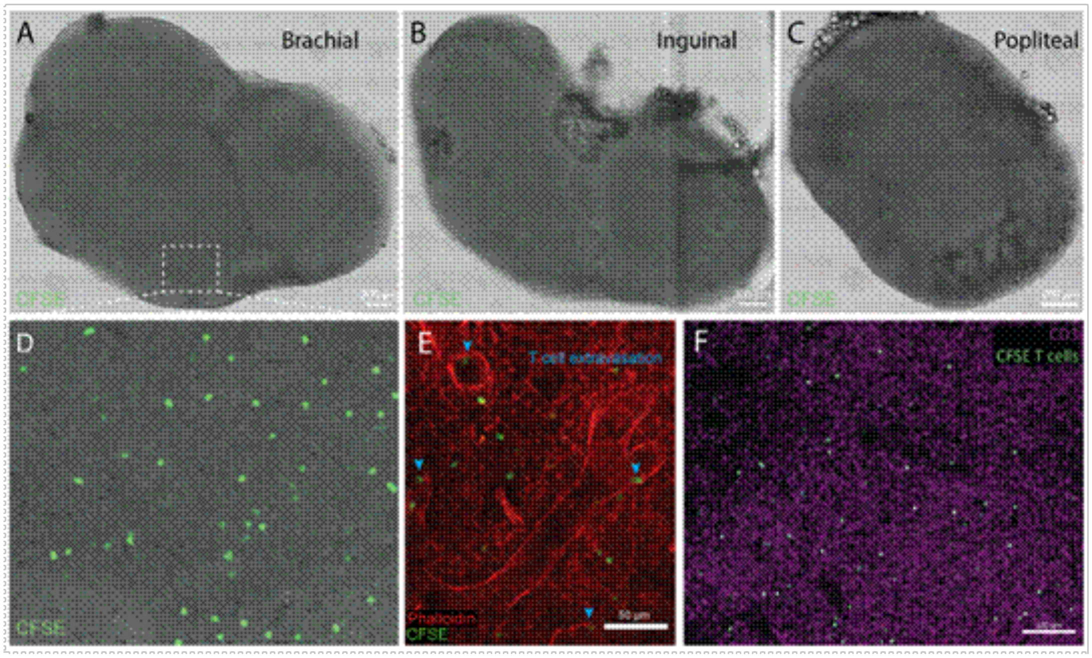


**Figure S3**: Following adoptive primary T lymphocyte transfer, CFSE-labeled cells are visible in A) brachial B) inguinal, and C) popliteal lymph nodes. D) Zoomed inset panel of a brachial lymph node shows homogenous distribution of cells. Counterstains with E) phalloidin and F) anti-CD3 antibody show interaction with local HEVs and normal lobule behavior. Dose response to L-selectin FAB.

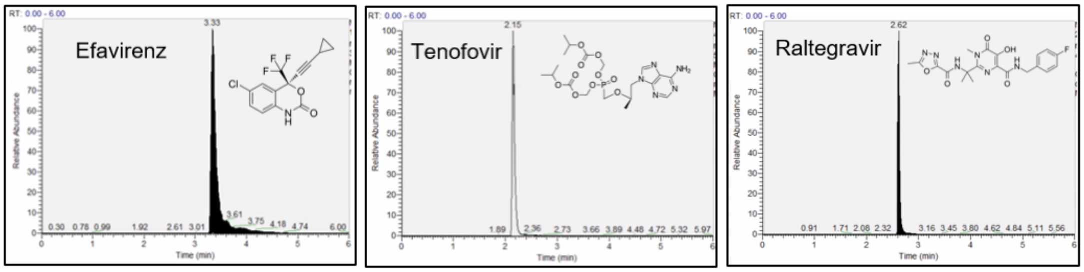


**Figure S4:** ARV species in this study are reliably extracted and detected at the following retention times.

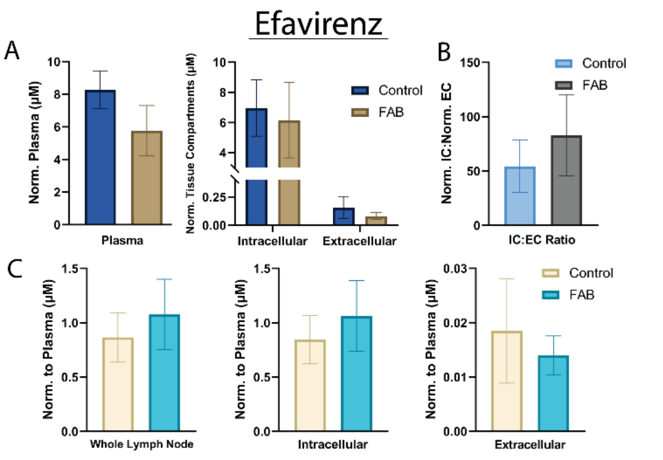


**Figure S5:** Efavirenz FAB results. (n=6)

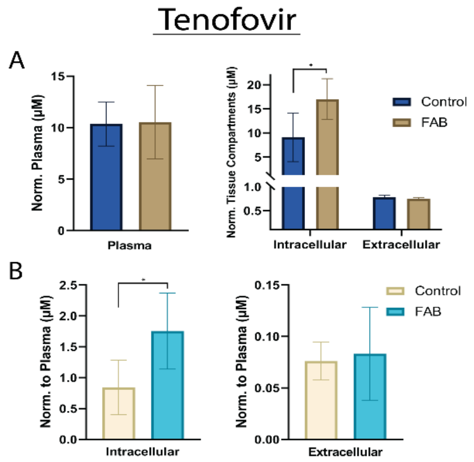


**Figure S6:** Tenofovir FAB results. (n=6)

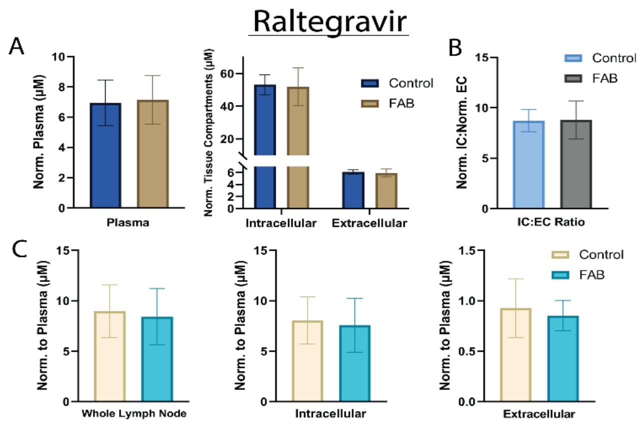


**Figure S7:** Raltegravir FAB results. (n=6)

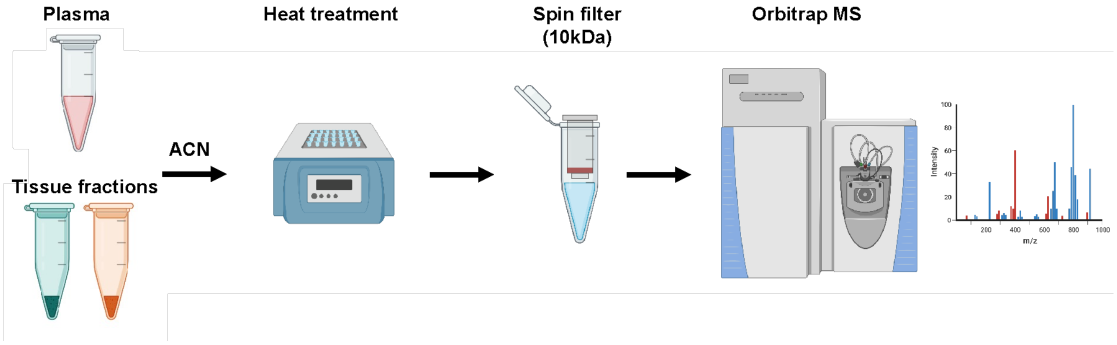


**Figure S8:** An overview of heat treatment processing of compartmentalized LN and plasma fractions for ARV extraction and MS analysis. Samples were treated with acidic acetonitrile and placed on a rocking heat block at 50°C and 350 rpm for 45 minutes. Samples were allowed to briefly cool before being vortexed and spun down to account for evaporation. Each sample was filtered through a 10 kDa filtered microcentrifuge tube. Filtrate was collected and submitted for MS analysis. Created using Biorender.com

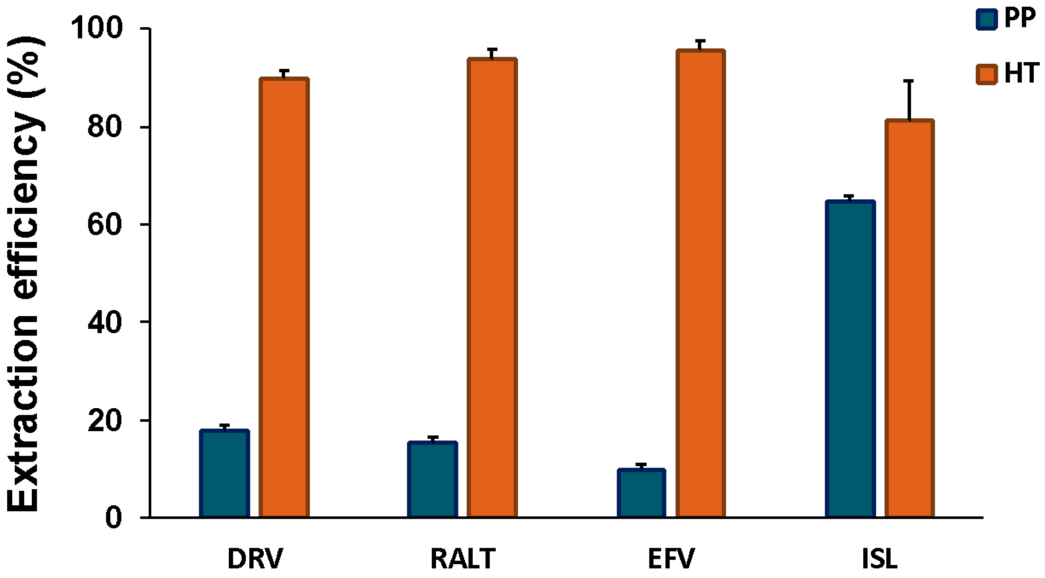


**Figure S9:** Determination of ARV extraction efficiency between protein precipitation and heat treatment extraction methods. Heat treatment demonstrated greater extraction efficiency across all ARVs in this study.

### COMPUTATIONAL SUPPLEMENTAL RESULTS AND FIGURES

#### Sensitivity Analysis

##### 1. Sensitivity Analysis of Peak Intracellular Tenofovir Concentration in the Lymph Node under FAB Condition:

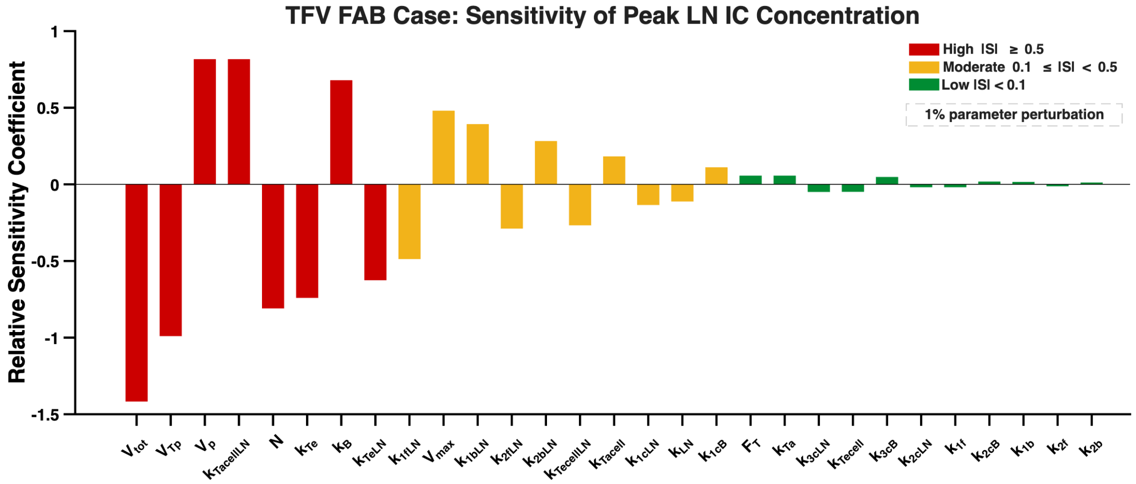


**Figure S10:** Sensitivity analysis of peak intracellular (IC) Tenofovir (TFV) concentration in the lymph node (LN) under FAB condition. Relative sensitivity coefficients were calculated using a 1% one-at-a-time parameter perturbation. Parameters were classified as highly sensitive (red, |S|≥0.5), moderately sensitive (yellow, 0.1≤|S|<0.5), or weakly sensitive (green, |S|<0.1 ).

##### 2. Sensitivity Analysis of Peak Intracellular Efavirenz Concentration in the Lymph Node under FAB Condition:

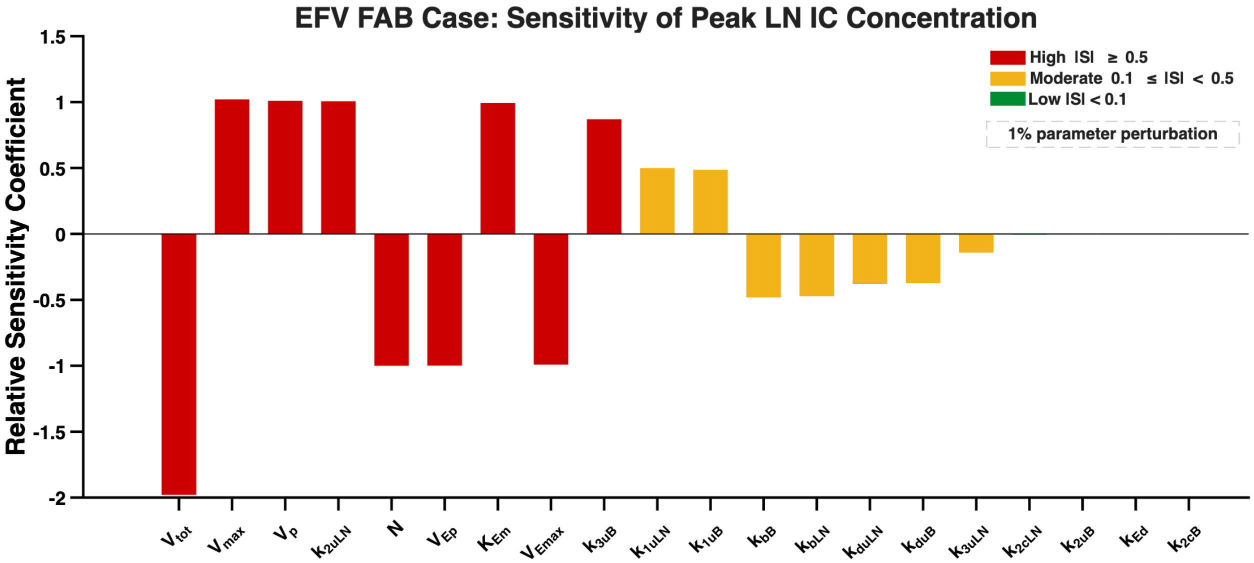


**Figure S11:** Relative sensitivity coefficients for the peak intracellular (IC) efavirenz (EFV) concentration in the lymph node (LN) under the FAB condition. Relative sensitivity coefficients were calculated using a 1% one-at-a-time parameter perturbation. Parameters were classified as highly sensitive (red, |S|≥0.5), moderately sensitive (yellow, 0.1≤|S|<0.5), or weakly sensitive (green, |S|<0.1 ).

#### 3. Sensitivity Analysis of Peak Intracellular Raltegravir Concentration in the Lymph Node under FAB Condition:

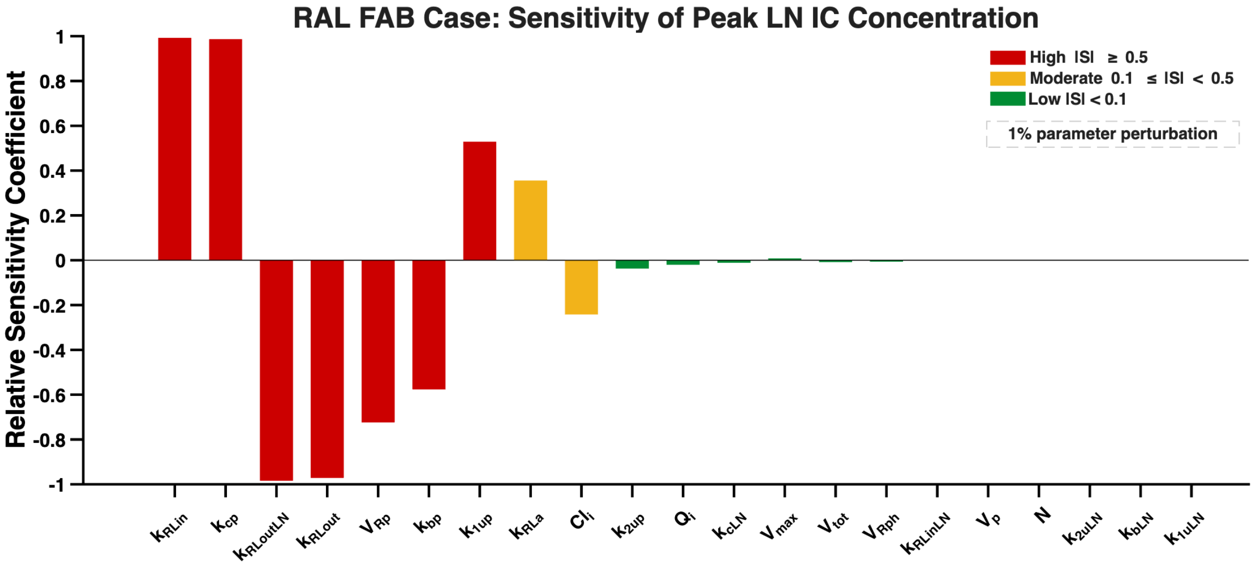


**Figure S12:** Relative sensitivity coefficients for the peak intracellular (IC) Raltegravir (RAL) concentration in the lymph node (LN) under the FAB condition. Relative sensitivity coefficients were calculated using a 1% one-at-a-time parameter perturbation. Parameters were classified as highly sensitive (red, $|S| \geq 0.5$), moderately sensitive (yellow, $0.1 \leq |S| < 0.5$), or weakly sensitive (green, $|S| < 0.1$ ).

### Antiretroviral Drug Concentration–Time Profiles in the Lymph Node

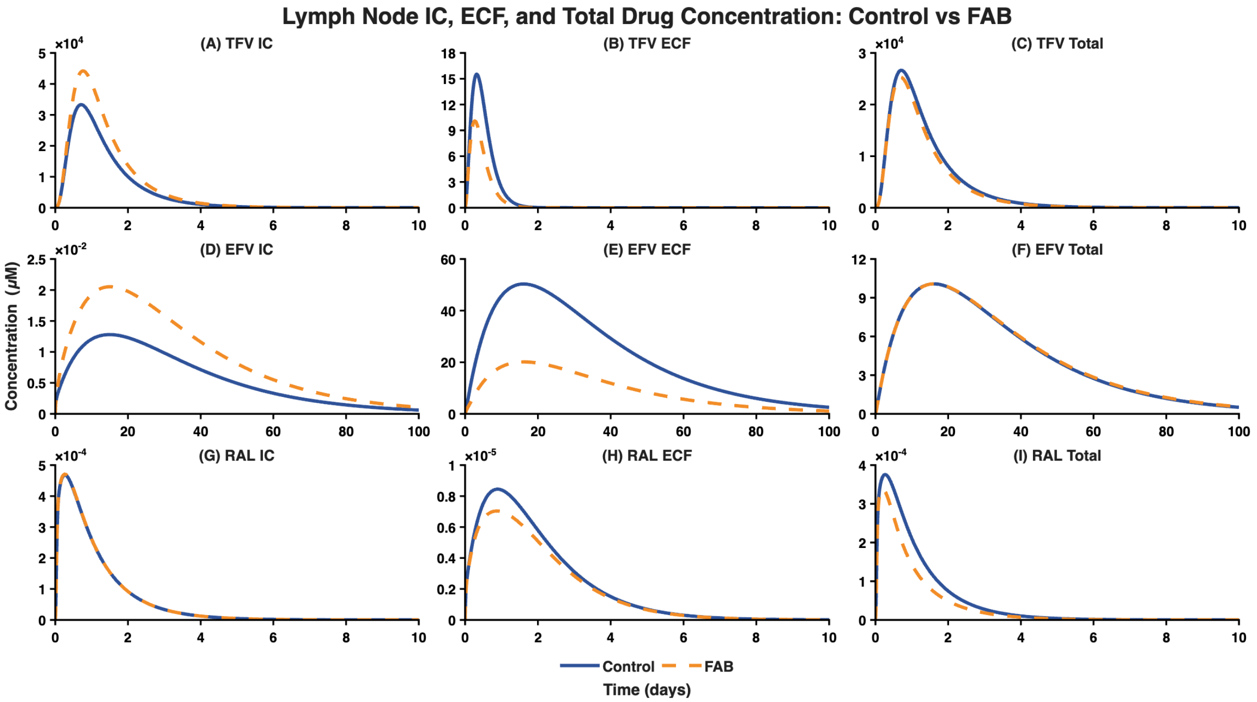


**Figure S13:** Lymph node intracellular (IC), extracellular fluid (ECF), and total antiretroviral drug concentrations under control and FAB condition. Simulated concentration–time profiles are shown for tenofovir (TFV; A–C), efavirenz (EFV; D–F), and raltegravir (RAL; G–I). For each drug, the first panel shows LN intracellular concentration, the second shows LN extracellular concentration, and the third shows total LN drug concentration. Solid blue lines represent control conditions, while dashed orange lines represent the FAB condition modeled through expansion of the LN extracellular fluid volume as a surrogate for reduced lymphocyte entry.

#### Drug Transport and Reaction Kinetics

The following equations describe the drug-specific transport and reaction kinetics used for EFV, TFV, and RAL. For each ARV, the model accounts for systemic absorption and elimination, plasma distribution, exchange between blood and LN compartments, and IC drug transport. Drug-specific processes, including reversible binding for EFV and RAL, IC phosphorylation for TFV, and RAL peripheral compartmental transport, were incorporated according to the pharmacological properties of each compound. The governing equations for each drug are provided below. All drug transport and reaction kinetic equations were implemented and solved in MATLAB R2022a using the built-in stiff ordinary differential equation solver `ode15s`.

#### Differential Equations of Efavirenz

##### Gut

$$\frac{dE_d}{dt} = -k_{Ed} \times E_d \quad \text{(Eq. S1)}$$

##### Blood

$$\frac{dE_{bB}}{dt} = k_{bB} \times E_{uB} - k_{1uB} \times E_{bB} - k_{dbB} \times E_{bB} \quad \text{(Eq. S2)}$$

$$\frac{dE_{uB}}{dt} = \frac{k_{Ed} \times E_d}{V_{EP}} - k_{bB} \times E_{uB} + k_{1uB} \times E_{bB} - k_{2uB} \times E_{uB} + \frac{V_{Emax} \times E_{cB}}{K_{Em} + E_{cB}} - k_{3uB} \times E_{uB} + k_{3uLN} \times E_{uLN} \times N \times \frac{V_{ex}}{V_p} - k_{duB} \times E_{uB} \quad \text{(Eq. S3)}$$

$$\frac{dE_{cB}}{dt} = k_{2uB} \times E_{uB} - \frac{V_{max} \times E_{cB}}{K_{Em} + E_{cB}} - k_{2cB} \times E_{cB} + k_{2cLN} \times E_{cLN} - k_{dcB} \times E_{cB} \quad \text{(Eq. S4)}$$

##### Lymph Node

$$\frac{dE_{bLN}}{dt} = k_{bLN} \times E_{uLN} - k_{1uLN} \times E_{bLN} - k_{dbLN} \times E_{bLN} - r_{inc} \times \left(1 - \frac{V_{ex}}{V_{max}}\right) \times E_{bLN} \quad \text{(Eq. S5)}$$

$$\frac{dE_{uLN}}{dt} = -k_{bLN} \times E_{uLN} + k_{1uLN} \times E_{bLN} - k_{2uLN} \times E_{uLN} + \frac{V_{Emax} \times E_{cLN}}{K_{Em} + E_{cLN}} \times \frac{V_{ic}}{V_{ex}} - k_{3uLN} \times E_{uLN} + k_{3uB} \times E_{uB} \times \frac{V_p}{V_{ex}} \times \frac{1}{N} - k_{duLN} \times E_{uLN} - r_{inc} \times \left(1 - \frac{V_{ex}}{V_{max}}\right) \times E_{uLN} \quad \text{(Eq. S6)}$$

$$\frac{dE_{cLN}}{dt} = k_{2uLN} \times E_{uLN} \times \frac{V_{ex}}{V_{ic}} - \frac{V_{Emax} \times E_{cLN}}{K_{Em} + E_{cLN}} - k_{2cLN} \times E_{cLN} + k_{2cB} \times E_{cB} - k_{dcLN} \times E_{cLN} + r_{inc} \times \left(1 - \frac{V_{ex}}{V_{max}}\right) \times \frac{V_{ex}}{V_{ic}} \times E_{cLN} \quad \text{(Eq. S7)}$$

**Differential Equations of Tenofovir**

**Gut**

$\frac{dT}{dt} = -kT_a \times FT \times Td$ (Eq. S8)

**Blood**

$\frac{dT_p}{dt} = \frac{(kT_a . \times FT \times Td)}{V_{Tp}} - kT_e \times T_p - k_B \times T_p + k_{LN} \times N \times \frac{V_{ex}}{V_p} \times T_{eLN}$ (Eq. S9)

$\frac{dT_c}{dt} = k_{Tacell} \times T_p - k_{Tecell} \times T_c - k_{1f} \times T_c + k_{1b} \times T_{cmp} - k_{1cB} \times T_c +$
$k_{1cLN} \times T_{cLN}$ (Eq. S10)

$\frac{dT_{cmp}}{dt} = k_{1f} \times T_c - k_{1b} \times T_{cmp} + k_{2b} \times T_{cdp} - k_{2f} \times T_{cmp} - k_{Tecell} \times T_{cmp} -$
$k_{2cB} \times T_{cmp} + k_{2cLN} \times T_{cmpLN}$ (Eq. S11)

$\frac{dT_{cdp}}{dt} = k_{2f} \times T_{cmp} - k_{2b} \times T_{cdp} - k_{Tecell} \times T_{cdp} - k_{3cB} \times T_{cdp} + k_{3cLN} \times T_{cdpLN}$ (Eq. S12)

**Lymph Node**

$\frac{dT_{eLN}}{dt} = -kT_{eLN} \times T_{eLN} + k_B \times T_p \times \frac{V_p}{V_{ex}} . \frac{1}{N} - k_{LN} \times T_{eLN} - r_{inc} \times \left(1 - \frac{V_{ex}}{V_{max}}\right) \times T_{eLN}$
(Eq. S13)

$\frac{dT_{cLN}}{dt} = k_{TacellLN} \times T_{eLN} \times \left(\frac{V_{ex}}{V_{ic}}\right) - k_{TecellLN} \times T_{cLN} - k_{1fLN} \times T_{cLN} + k_{1bLN} \times T_{cmpLN} +$
$k_{1cB} \times T_c - k_{1cLN} \times T_{cLN} + \left(\frac{\left(r_{inc} \times \left(1 - \frac{V_{ex}}{V_{max}}\right) \times V_{ex}\right)}{V_{ic}}\right) \times T_{cLN}$ (Eq. S14)

$\frac{dT_{cmpLN}}{dt} = k_{1fLN} \times T_{cLN} - k_{1bLN} \times T_{cmpLN} + k_{2bLN} \times T_{cdpLN} - k_{2fLN} \times T_{cmpLN} -$
$k_{TecellLN} \times T_{cmpLN} + k_{2cB} \times T_{cmp} - k_{2cLN} \times T_{cmpLN} + \left(\frac{\left(r_{inc} \times \left(1 - \frac{V_{ex}}{V_{max}}\right) \times V_{ex}\right)}{V_{ic}}\right) \times T_{cmpLN}$
(Eq. S15)

$$\frac{dT_{cdpLN}}{dt} = k_{2fLN} \times T_{cmpLN} - k_{2bLN} \times T_{cdpLN} - k_{TecellLN} \times T_{cdpLN} + k_{3cB} \times T_{cdp} - k_{3cLN} \times T_{cdpLN} + \left(\frac{\left(r_{inc} \times \left(1 - \frac{(V_{ex})}{V_{max}}\right) \times V_{ex}\right)}{V_{ic}}\right) \times T_{cdpLN} \quad \text{(Eq. S16)}$$

**Differential Equations of Raltegravir**

**Gut**

$$\frac{dRL}{dt} = -k_{RLa} \times RL \quad \text{(Eq. S17)}$$

**Blood**

$$\frac{dRL_{bp}}{dt} = -k_{1up} \times RL_{bp} + k_{bp} \times RL_{up} - k_{ebp} \times RL_{bp} \quad \text{(Eq. S18)}$$

$$\frac{dRL_{up}}{dt} = k_{RLa} \times RL / V_{ep} - k_{bp} \times RL_{up} + k_{1up} \times RL_{bp} - k_{RLin} \times RL_{up} + k_{RLout} \times RL_{c} - k_{2up} \times RL_{up} + k_{2uLN} \times N \times R_{LuLN} \times \frac{V_{ex}}{V_p} - \left(\frac{Cli}{V_1}\right) \times RL_{up} - \left(\frac{Qi}{V_1}\right) \times RL_{up} + \left(\frac{Qi}{V_1}\right) \times RL_{ph} \quad \text{(Eq. S19)}$$

$$\frac{dRL_{ph}}{dt} = \left(\frac{Qi}{V_2}\right) \times RL_{up} - \left(\frac{Qi}{V_2}\right) \times RL_{ph} \quad \text{(Eq. S20)}$$

$$\frac{dRL_{c}}{dt} = k_{RLin} \times RL_{up} - k_{RLout} \times RL_{c} - k_{cp} \times RL_{c} + k_{cLN} \times RL_{cLN} - k_{ec} \times RL_{c} \quad \text{(Eq. S21)}$$

**Lymph Node**

$$\frac{dRL_{bLN}}{dt} = -k_{1uLN} \times RL_{bLN} + k_{bLN} \times RL_{uLN} - k_{ebLN} \times RL_{bLN} - r_{inc} \times \left(1 - \frac{V_{ex}}{V_{max}}\right) \times RL_{bLN} \quad \text{(Eq. S22)}$$

$$\frac{dRL_{uLN}}{dt} = k_{1uLN} \times RL_{bLN} - k_{bLN} \times RL_{uLN} - k_{RLinLN} \times RL_{uLN} + k_{RLoutLN} \times RL_{cLN} \times \frac{V_{ic}}{V_{ex}} + k_{2up} \times RL_{up} \times \frac{V_p}{V_{ex}} \times \frac{1}{N} - k_{2uLN} \times RL_{uLN} - k_{euLN} \times RL_{uLN} - r_{inc} \times \left(1 - \frac{V_{ex}}{V_{max}}\right) \times R_{LuLN} \quad \text{(Eq. S23)}$$

$$\frac{dRL_{cLN}}{dt} = k_{RLinLN} \times RL_{uLN} \times \frac{V_{ex}}{V_{ic}} - k_{RLoutLN} \times RL_{cLN} + k_{cp} \times RL_{c} - k_{cLN} \times RL_{cLN} - k_{ecLN} \times RL_{cLN} + r_{inc} \times \left(1 - \frac{V_{ex}}{V_{max}}\right) \times \frac{V_{ex}}{V_{ic}} \times RL_{cLN} \quad \text{(Eq. S24)}$$

#### Parameter Values

**Table S2.** PBPK Model Parameters for Efavirenz (EFV)

| Sr. No | Parameters | Parameter name | Value | Unit |
|---|---|---|---|---|
| 1. | $E_d$ | Efavirenz dose | 600 | mg day$^{-1}$ |
| 2. | $k_{Ed}$ | Drug absorption rate constant from gut to plasma | 0.20 × 24 | day$^{-1}$ |
| 3. | $k_{1uB}$ | Unbinding rate in blood plasma | 10 | day$^{-1}$ |
| 4. | $k_{bB}$ | Binding rate in blood plasma | 50 × $k_{1uB}$ | day$^{-1}$ |
| 5. | $k_{1uLN}$ | Unbinding rate in LN ECF | 1 | day$^{-1}$ |
| 6. | $k_{bLN}$ | Binding rate in LN ECF | 50 × $k_{1uLN}$ | day$^{-1}$ |
| 7. | $V_{Ep}$ | Effective volume distribution in plasma | 100 × $10^3$ | mL |
| 8. | $V_p$ | Plasma volume | 3 × $10^3$ | mL |
| 9. | $k_{Ein}$ | Rate constant for transfer of EFV from Plasma to PBMC (blood cells) | 0.32 × 24 | day$^{-1}$ |
| 10. | $k_{2uB}$ | Plasma-to-blood-cell transfer rate | $k_{Ein}$ | day$^{-1}$ |
| 11. | $k_{2uLN}$ | LN ECF-to-cell uptake rate | $k_{2uB}$ | day$^{-1}$ |
| 12. | $k_{Ee}$ | Elimination rate constant for EFV from plasma | 0.10 × 24 | day$^{-1}$ |
| 13. | $k_{duLN}$ | Unbound LN ECF decay rate | $k_{Ee}$ | day$^{-1}$ |
| 14. | $k_{dbLN}$ | Bound LN ECF decay rate | 0 | day$^{-1}$ |
| 15. | $k_{duB}$ | Unbound plasma decay rate | $k_{Ee}$ | day$^{-1}$ |
| 16. | $k_{dbB}$ | Bound plasma decay rate | 0 | day$^{-1}$ |
| 17. | $k_{dcB}$ | Blood intracellular decay rate | 0 | day$^{-1}$ |
| 18. | $k_{dcLN}$ | LN intracellular decay rate | 0 | day$^{-1}$ |
| 19. | $N$ | Number of lymph nodes | 600 | — |
| 20. | $V_{me}$ | Maximum rate for transfer of EFV from PBMC to plasma | (4400 × $10^{-6}$) × 24 | mg mL day$^{-1}$ |
| 21. | $K_{me}$ | Average Michaelis Menten constant for transfer of EFV from PBMC to Plasma | 710 × $10^{-6}$ | mg mL$^{-1}$ |
| 22. | $k_{2cB}$ | Blood-cell to LN-cell transfer rate | 1 | day$^{-1}$ |
| 23. | $k_{2cLN}$ | LN-cell to blood-cell transfer rate | $k_{2cB}$ | day$^{-1}$ |
| 24. | $k_{3uB}$ | Blood plasma-to-LN ECF transfer rate | $k_{2cB}$ | day$^{-1}$ |
| 25. | $k_{3\_uLN}$ | LN ECF-to-blood plasma transfer rate | $k_{3uB}$ | day$^{-1}$ |
| 26. | $V_{max}$ | Maximum LN expansion | 0.5 | mL |
| 27. | $V_{tot}$ | Total LN volume | 1 | mL |

**Table S3.** PBPK Model Parameters for Tenofovir (TFV)

| Sr. No | Parameter | Parameter name | Value | Unit |
|---|---|---|---|---|
| 1. | $T_d$ | Tenofovir dose | 300 | mg day$^{-1}$ |
| 2. | $k_{Ta}$ | Drug absorption rate constant from gut to plasma | 19.72 | day$^{-1}$ |
| 3. | $F_T$ | Bioavailability of TFV | 0.39 | — |
| 4. | $V_{Tp}$ | Volume of Distribution in blood for TFV | 87360 | mL |
| 5. | $k_{Te}$ | Elimination rate constant from plasma | 4.74 | day$^{-1}$ |
| 6. | $k_B$ | Plasma-to-LN ECF transfer rate | $k_{1cB}$ | day$^{-1}$ |
| 7. | $k_{LN}$ | LN ECF-to-cell uptake rate | $k_B$ | day$^{-1}$ |
| 8. | $k_{Tacell}$ | Plasma-to-blood-cell transfer rate | 24000 | day$^{-1}$ |
| 9. | $f_{BT}$ | TDF protein binding factor | 0.07 | — |
| 10. | $H_T$ | Partition coefficient | 1800 | — |
| 11. | $k_{Tecell}$ | Blood intracellular elimination rate | 1.1 | day$^{-1}$ |
| 12. | $k_{1f}$ | Forward monophosphate conversion rate | 9.6 | day$^{-1}$ |
| 13. | $k_{1b}$ | LN reverse monophosphate conversion rate | 30.3 | day$^{-1}$ |
| 14. | $k_{1cB}$ | Blood-to-LN TFV intracellular transfer rate | 1 | day$^{-1}$ |
| 15. | $k_{1cLN}$ | LN-to-blood TFV intracellular transfer rate | $k_{1cB}$ | day$^{-1}$ |
| 16. | $k_{2b}$ | Reverse diphosphate conversion rate | 95.5 | day$^{-1}$ |
| 17. | $k_{2f}$ | Forward diphosphate conversion rate | 270.7 | day$^{-1}$ |
| 18. | $k_{2cB}$ | Blood-to-LN TFV monophosphate intracellular transfer rate | $k_{1cB}$ | day$^{-1}$ |
| 19. | $k_{2cLN}$ | LN-to-Blood TFV diphosphate intracellular transfer rate | $k_{1cB}$ | day$^{-1}$ |
| 20. | $k_{3cB}$ | Blood-to-LN TFV diphosphate intracellular transfer rate | $k_{1cB}$ | day$^{-1}$ |
| 21. | $k_{3cLN}$ | LN-to-Blood TFV monophosphate intracellular transfer rate | $k_{1cB}$ | day$^{-1}$ |
| 22. | $k_{TeLN}$ | LN extracellular elimination rate | $k_{Te}$ | day$^{-1}$ |
| 23. | $k_{TacellLN}$ | LN ECF-to-LN-cell transfer rate | 24000 | day$^{-1}$ |
| 24. | $f_{BTLN}$ | TDF protein binding factor | $f_{BT}$ | — |
| 25. | $H_{TLN}$ | Partition coefficient | $H_T$ | — |
| 26. | $k_{TecellLN}$ | Partition coefficient | $k_{Tecell}$ | day$^{-1}$ |
| 27. | $k_{1fLN}$ | LN forward monophosphate conversion rate | $k_{1f}$ | day$^{-1}$ |
| 28. | $k_{1bLN}$ | LN forward monophosphate conversion rate | $k_{1b}$ | day$^{-1}$ |
| 29. | $k_{2bLN}$ | LN reverse diphosphate conversion rate | $k_{2b}$ | day$^{-1}$ |
| 30. | $k_{2fLN}$ | LN forward diphosphate conversion rate | $k_{2f}$ | day$^{-1}$ |

**Table S4.** PBPK Model Parameters for Raltegravir (RAL)

| **Sr. No** | **Parameter** | **Parameter name** | **Value** | **Unit** |
|---|---|---|---|---|
| 1 | RL | Raltegravir dose | 400 mg (twice daily) | mg day$^{-1}$ |
| 2 | $k_{RLa}$ | Drug absorption rate constant (gut to plasma) | 0.21 × 24 | day$^{-1}$ |
| 3 | $k_{1up}$ | Unbinding rate in blood plasma | 10 | day$^{-1}$ |
| 4 | $k_{bp}$ | Binding rate in blood plasma | 5 × $k_{1up}$ | day$^{-1}$ |
| 5 | $k_{1uLN}$ | Unbinding rate in LN ECF | 1 | day$^{-1}$ |
| 6 | $k_{bLN}$ | Binding rate in LN ECF | 5 × $k_{1uLN}$ | day$^{-1}$ |
| 7 | $k_{RLin}$ | Rate constant for RL transfer (plasma to blood cells) | 0.1956 × 24 | day$^{-1}$ |
| 8 | $k_{RLout}$ | Rate constant for RL transfer (blood cells to plasma) | 3.69 × 24 | day$^{-1}$ |
| 9 | $k_{RLinLN}$ | Rate constant for RL transfer (LN ECF to LN cells) | $k_{RLin}$ | day$^{-1}$ |
| 10 | $k_{RLoutLN}$ | Rate constant for RL transfer (LN cells to LN ECF) | $k_{RLout}$ | day$^{-1}$ |
| 11 | $k_{cp}$ | Blood-cell to LN-cell transfer rate | 1 | day$^{-1}$ |
| 12 | $k_{cLN}$ | LN-cell to blood-cell transfer rate | $k_{cp}$ | day$^{-1}$ |
| 13 | $k_{2uLN}$ | LN ECF-to-cell uptake rate | $k_{cp}$ | day$^{-1}$ |
| 14 | $k_{2up}$ | Plasma-to-blood-cell transfer rate | $k_{2uLN}$ | day$^{-1}$ |
| 15 | $Cl_i$ | Intrinsic clearance rate | 60.2 × 24 | L/day |
| 16 | $Q_i$ | Intracellular flow / perfusion rate | 8.5 × 24 | L/day |
| 17 | $k_{ebp}$ | Bound plasma decay rate | 0 | day$^{-1}$ |
| 18 | $k_{ec}$ | Blood intracellular decay rate | 0 | day$^{-1}$ |
| 19 | $k_{ebLN}$ | Bound LN ECF decay rate | 0 | day$^{-1}$ |
| 20 | $V_{Rp}$ | Volume of distribution for central compartment i.e. plasma | 223 | L |
| 21 | $V_{Rph}$ | Volume of distribution for peripheral compartment | 113 | L |
| 22 | $k_{ecLN}$ | LN intracellular decay rate | 0 | day$^{-1}$ |
